# Imbalance in gut microbial interactions as a marker of health and disease

**DOI:** 10.1101/2025.04.30.651474

**Authors:** Roberto Corral López, Juan A. Bonachela, Maria Gloria Dominguez-Bello, Michael Manhart, Simon A. Levin, Martin J. Blaser, Miguel A. Muñoz

**Affiliations:** Departamento de Electromagnetismo y Física de la Materia. Universidad de Granada. Granada, Spain; Instituto Carlos I de Física Teórica. Universidad de Granada. Granada, Spain; Department of Ecology, Evolution, and Natural Resources, Rutgers University, New Brunswick, NJ, USA; Department of Biochemistry and Microbiology, Rutgers University, New Brunswick, NJ, USA; Center for Advanced Biotechnology and Medicine, Rutgers University, Piscataway, NJ, USA; Department of Biochemistry and Molecular Biology, Rutgers RWJ Medical School, Piscataway, NJ, USA; Department of Ecology and Evolutionary Biology, Princeton University, Princeton, NJ, USA

## Abstract

Imbalances in the human gut microbiome (dysbioses) are linked to multiple diseases but remain poorly understood. Current biomarkers to identify dysbiosis are inconsistent and fail to capture the ecological mechanisms differentiating healthy from diseased microbiomes. We propose a general dysbiosis biomarker, inspired by phenomenology observed in a gut-microbiome theoretical model introduced here. The emergent communities show complex interaction networks and two distinct collective states, corresponding to healthy and dysbiotic microbiomes. Our robust metric for dysbiosis, by quantifying the balance between cooperation and competition, differentiates these states in both simulated and real datasets across diverse diseases. Moreover, it reveals that dysbiosis results from a shift toward greater cooperation in the community. Our metric further correlates with disease progression, highlighting its potential as a diagnostic tool.

---

The human gut microbiome is a dynamic and complex system that plays a crucial role in maintaining health through its interactions with the host and environment (*1–6*). Innovations in profiling techniques over the past two decades, have greatly improved our ability to quantify both microbiome composition and functional capabilities (*7–9*). These studies have linked imbalances in the gut microbiome, or dysbioses, to numerous diseases, including obesity, diabetes, inflammatory bowel disease (IBD), *Clostridioides difficile* infection (CDI), irritable bowel syndrome (IBS), colorectal cancer (CRC), liver disease, pancreatic disorders, psoriatic arthritis, and celiac disease, among others (*1, 2, 7, 10–16*). Consequently, novel treatments such as fecal microbiota transplantation, diet-based interventions, and probiotics (*17–19*), aim to restore microbial balance and reverse dysbiosis by altering community composition to eliminate such imbalances. However, these treatments show highly variable and unpredictable success rates, which slows clinical implementation. A key reason for this unpredictability is the limited understanding of the mechanisms underlying gut microbiome dynamics and structure. Particularly, the metabolic and ecological interactions among microorganisms, and their relationships with the host and environment, remain poorly understood (*1, 14, 20, 21*). This knowledge gap has hindered the identification of robust microbiome biomarkers that correlate with, and accurately differentiate, health and disease. Our work seeks to address both of these fundamental challenges.

## 1 The search for a universal indicator of dysbiosis

Dysbiosis is a broad concept that encompasses various compositional and functional attributes across diseases, which hinders its unambiguous characterization (*12*). Early efforts focused on identifying single bacterial species or functions whose altered levels could correlate with a particular disease (*10, 12, 22, 23*). However, it soon became clear that dysbiosis is a collective state, typically affecting multiple species and functions, which can only be captured by systemic, whole-community approaches (*16, 24*). A common claim across studies and conditions is that disease correlates with reduced microbial diversity, often suggested as a universal marker of dysbiosis (*7,13,15,25–27*). We re-examined empirical data from several diseases, and our analysis reveals a more nuanced, disease-specific pattern (Fig. 1), in line with recent studies (*12, 16, 24, 28*). Specifically, while IBD and CDI (Fig. 1A-B) consistently show reduced diversity compared to healthy microbiomes (*16, 29–31*), conditions like IBS and CRC (Fig. 1C-D) yield mixed results, with decreases, increases, or no statistically significant change in diversity (*16, 32–37*). These findings clearly refute that diversity universally declines in diseased states, and raise the fundamental question of whether there are conserved mechanisms driving dysbiosis, which therefore would enable the identification of robust disease indicators.

**Figure 1:**
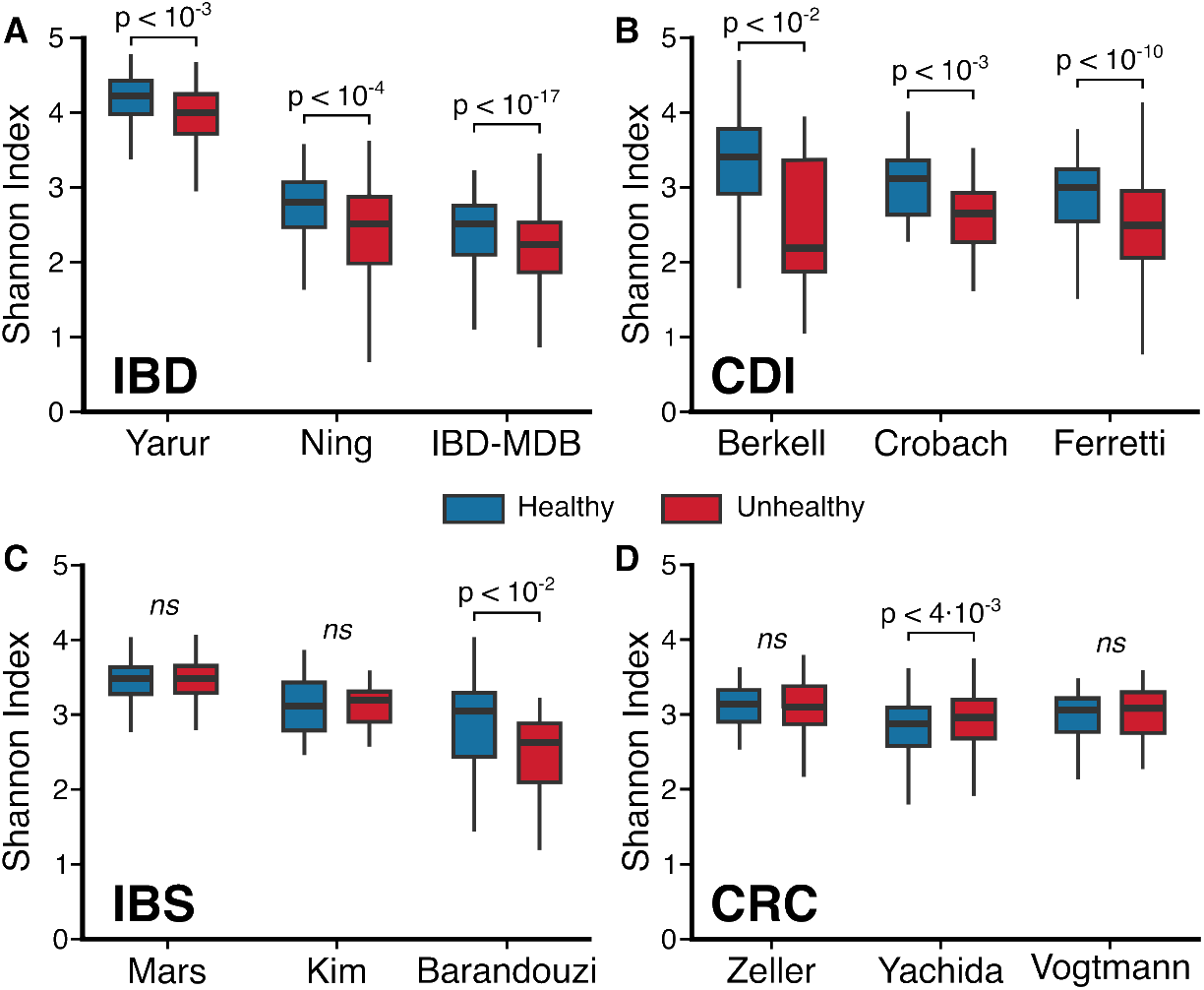
*α*-diversity in healthy and unhealthy individuals. Comparison of the Shannon index (see Methods) obtained using existing metagenomic analyses from healthy individuals, in blue, whereas the red color represents individuals affected by : (**A**) inflammatory bowel disease (IBD), using data from Yarur et al., Ning et al., and the Human Microbiome Project (*38–40*); (**B**) *C. difficile* infection (CDI), using data from Berkell et al., Crobach et al., and Ferreti et al. (*41–43*); (**C**) irritable bowel syndrome (IBS), using data from Mars et al., Kim et al., and Barandouzi et al. (*44–46*); and (**D**) colorectal cancer (CRC), using data from Zeller et al., Yachida et al., and Vogtmann et al. (*37, 47–49*). A significant decrease in diversity is evident for IBD and CDI, but not for IBS or CRC (*ns* = not significant).

The dichotomy between health and disease has fuelled the extensively explored hypothesis that dysbiosis generally emerges from shifts between alternative stable states, that is, between distinct taxonomic or functional configurations of the gut microbiome that can occur for identical conditions (*50–62*). However, this idea remains under debate due to the varied methodologies used to define and measure such states, the context-dependent nature of the microbiome, and the challenges of controlling external factors such as diet, medication, and lifestyle (*7, 10, 63*). All these factors influence gut composition and function, complicating the identification of stable states and consistent disease markers.

The challenges above, however, do not apply to theoretical models, where definitions and environmental factors can be fine-tuned (*21, 63–67*). For example, models have identified microbial taxa capable of inhibiting *C. difficile* (CD), a prediction subsequently validated experimentally to confirm that the suggested taxa indeed conferred CD colonization resistance (*68*). As another example, machine learning models have systematically tested competing hypotheses regarding the success of FMT (*69*). Despite these advances, few models designed to study gut microbiome dynamics predict or can even generate alternative stable states, limiting our understanding of their emergence and fundamental properties (*64,70*). Therefore, there is need for novel theoretical models that can capture gut microbiome dynamics and also reveal the ecological mechanisms driving the formation of diverse collective states.

## 2 A new model for the dynamics of the gut microbiome

We introduce here a mathematical model specifically designed to explore the ecological dynamics of the gut microbiome (Fig.2A and Supplementary Text). Our consumer-resource model captures bacterial and nutrient dynamics and incorporates the possibility of cross-feeding as new metabolites are produced by different bacterial strains. Nutrients are introduced at a rate *h* (representing host diet), new bacterial species immigrate at a rate *U*, and bacteria, nutrients, and metabolites are removed at a dilution rate *δ*, which determines intestinal time.

A key strength of our framework is its emphasis on the crucial role that metabolic pathways play in shaping gut communities (*71–79*). To this end, our model incorporates metabolic pathways that define all possible metabolic routes between nutrients and metabolites (hereon, substances), along with the metabolic preferences specific to each bacterial strain. While we explicitly represent only catabolic reactions for simplicity, anabolism is indirectly accounted for through a trade-off function linking enzymatic cost to energy invested in constructive metabolism. Thus, our model captures both the energy gain and enzymatic cost for each metabolic reaction.

To model the metabolic conversion of a substance *S*_*α*_ into another substance *S*_*β*_, we monitor 1) the free energy (*E*_*αβ*_ > 0) released during the process; and 2) the enzymatic cost (*C*_*αβ*_ > 0), reflecting the number and complexity of enzymes involved. Each bacterial type *i* is defined by a unique pathway matrix, 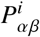 (binary matrix with the metabolic routes available to that bacterial type) and a coefficient *γ*_*i*_ that controls the fraction of free energy used for biomass growth. See Supplementary Text for a detailed description of the model, including the definition of species, nutrients, metabolites, the implementation of bacteria-specific metabolic preferences, pathways, trade-off function, and chosen parametrization. We examine the limitations associated with the modeling assumptions above in Discussion.

Our model successfully reproduces, without the need to rely on parameter fitting, well-known macroecological patterns of the gut microbiome that are observed empirically across multiple systems. Using a biologically plausible parametrization (see Supplementary Text), the 95% confidence interval obtained across replicates of the model captures the distributions of species abundance mean and fluctuations, Taylor’s law (i.e. relationship between mean and variance), species prevalence (the distribution of the fraction of samples in which a species is present), and the dependence of prevalence on mean abundance (Supplementary Fig.S1) (*80, 81*). Additionally, Supplementary Fig. S2 shows that our model replicates the functional redundancy observed in the gut microbiome, where multiple taxonomic compositions encode the same metabolic functions (*82–85*). In our framework, functions are represented by metabolites grouped into coarse-grained energy levels (see Methods (*86*)). These broader functional categories are akin to Clusters of Orthologous Genes (COG) or Kyoto Encyclopedia of Genes and Genomes (KEGG) classifications for empirical data (*87, 88*). Functional redundancy becomes apparent at sufficiently coarse-grained resolutions for both our model and observed data. All the results above thus validate our modeling approach.

## 3 Alternative states in the gut microbiome model may reflect health and disease

As shown in Figure 2B, the dynamics of our simulated gut microbiome community suggest two qualitatively distinct classes of states: one characterized by the rapid turnover of many bacterial types; and another one where only a few types dominate with quasi-stationary abundances (see definition in Methods (*86*)). Let us label these two classes of states as *healthy* and *dysbiotic*, respectively, for reasons that will become apparent in the comparison with real data below. Notably, these alternative states emerge consistently across a wide range of parameter values, which indicates that they result from the mechanisms encoded in the model rather than specific parametrizations (see Supplementary Fig. S3 and S4).

**Figure 2:**
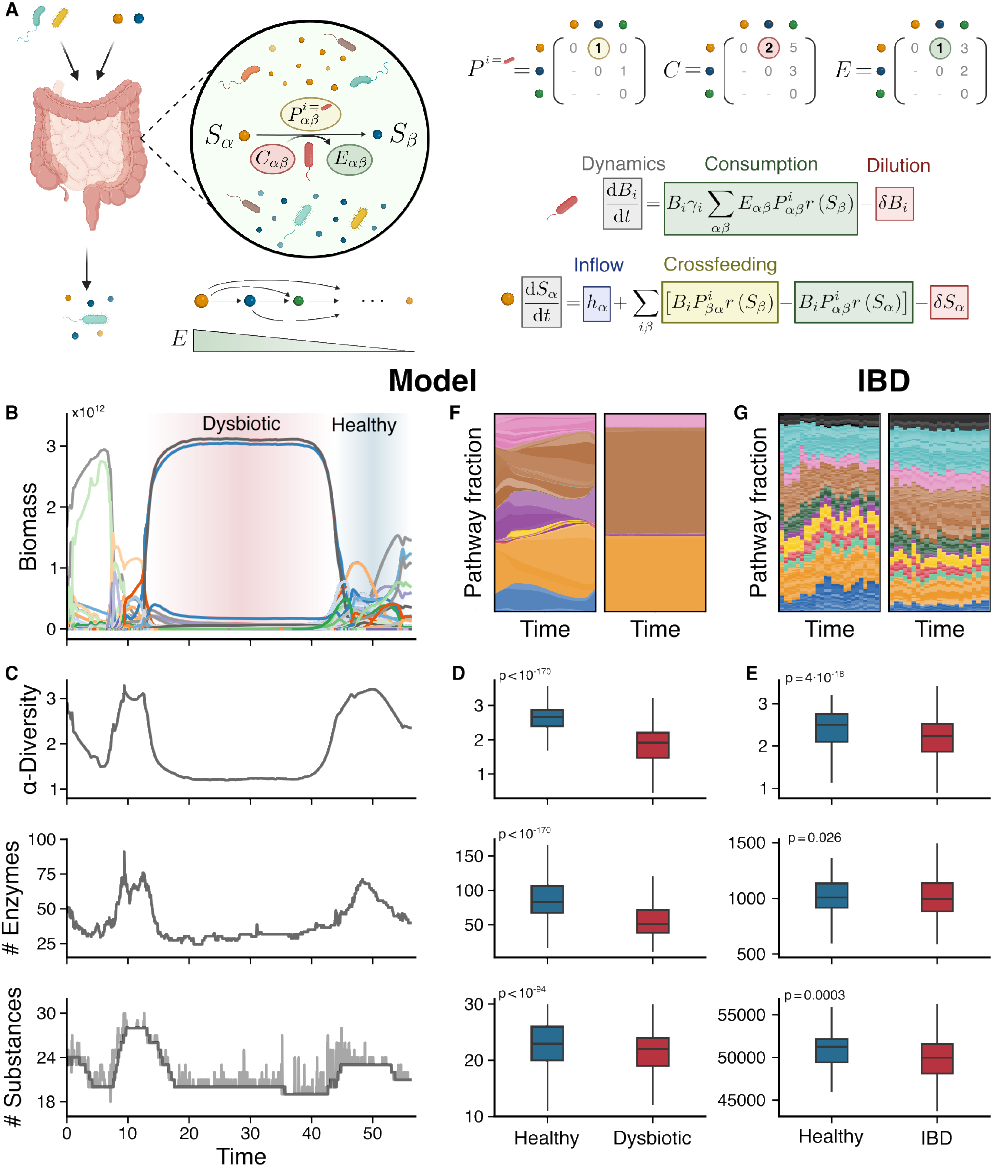
Gut microbiome trait-based consumer-resource model and the spontaneous emergence of alternative stable states. (**A**) Schematic representation of the model. The gut microbiome is conceptualized as living in a chemostat-like environment that includes both bacterial species and substances. Metabolic pathways linking any two substances, *S*_*α*_ and *S*_*β*_, are characterized by their enzymatic cost, *C*_*αβ*_, and the energy obtained from the conversion, *E*_*αβ*_. Each bacterial species can use specific pathways depending on its pathway matrix, *P*^*i*^, which determines the metabolic routes available to that species. The model considers only catabolic pathways, where *E*_*αβ*_ > 0. (**B**) Biomass (cells) over time for a realization of the model, illustrating the emergence of dysbiotic and healthy states. (**C**) For the realization highlighted in the larger panel B, calculated *α*-diversity, number of enzymes, and number of substances over time for a single realization of the model; dark grey represents rolling average, and light grey the actual values. (**D**) Comparison, for the same metrics in panel C, of healthy and dysbiotic states. (**E**) Corresponding metrics for healthy individuals and IBD patients. **(F)** Coarse-grained pathway fraction over time for a healthy (left) and dysbiotic (right) state in the model; colors represent different (coarse-grained) functions. **(G)** Similary plot for healthy/IBD, where colors represent (coarse-grain) functions, and shades the various paths able to perform the given function. See Methods for details (*86*).

Existing community-level theoretical models, such as the generalized Lotka-Volterra (GLV) (*89–91*) and consumer-resource models with random interactions (*92, 93*), have reported dynamics reminiscent of either one of our two states (often referred to as “chaotic fluctuations” and “stable fixed points” in the literature). However, these models do not capture multistability between the different states. The fact that our model captures diverse alternative states and dynamic transitions between them highlights the importance of explicitly incorporating metabolic pathways, and of letting the structure of the community emerge from the dynamic interaction in the system, elements overlooked in previous theoretical efforts. As shown in Fig. 2C-D, the dysbiotic state exhibits a significant reduction in *α*-diversity (measured by the Shannon Index), the number of enzymes (total enzymatic cost of all active pathways), and the number of secreted metabolites (substances with non-zero concentrations) compared to the healthy state; see Methods for definitions (*86*). These findings suggest that the microbial community in the dysbiotic state operates with greater efficiency than the community in the healthy state, as fewer bacterial types and metabolic pathways are required to form consortia able to convert a larger proportion of the available energy into microbial biomass (Supplementary Fig. S5).

Given the dichotomy between these two distinct states, reminiscent of the healthy vs dysbiotic gut microbiome, we also measured the biomarkers above in healthy individuals and patients with IBD. As Fig. 2E shows, the resulting pattern mirrors our model predictions, thereby supporting our designation of the two alternative states as *healthy* and *dysbiotic*. The parallelism of model states with real healthy/dysbiotic states is reinforced by the observation that, from a qualitative perspective, both in the model and in IBD the shift toward dysbiosis involves the loss of specific metabolic pathways, resulting in a disruption of functional capacity (that is, some functions may be impaired or entirely lost, while others become overrepresented, see Fig. 2F,G) (*94, 95*). Moreover, for our model as well as a range of diseases, *β*-diversity across healthy states was significantly lower than for dysbiotic states (Supplementary Fig. S6). These patterns, however, are subtle or inconsistent (see, for example, *β*-diversity for CRC, Supplementary Fig. S6D, or number of metabolites and diversity of secreted metabolites for IBS and CRC, Supplementary Fig. S7). Beyond this lack of consistency, these biomarkers merely reflect the consequences of the shift from one state to the other and therefore do not explain the mechanisms underlying the transitions, nor can be used as early-warning indicators. Thus, we further explored with our model the differences between healthy and dysbiotic states, aiming to develop a robust quantitative metric able to reliably distinguish between healthy and dysbiotic states and that can also reveal the mechanisms driving the emergence of these states and transitions.

## 4 The balance of community interactions is disrupted in dysbiotic states and disease

In our model, bacterial interactions are not assumed but rather emerge from competition for shared resources and cross-feeding. Community dynamics ultimately determine which substances (and, indirectly, which interactions) persist in the system. Therefore, our model enables a shift of focus from broad measures of species and functional diversity to the emergent, dynamic ecological network of effective interactions that collectively shape the resulting community (*64, 96, 97*).

There is currently no consensus on whether competition or cooperation primarily drive gut microbiota dynamics. While some empirical studies suggest that competitive interactions dominate due to their role in promoting stability (*21, 98, 99*), others emphasize the critical role of cross-feeding interactions in maintaining a healthy microbial community (*100–103*). Thus, the relative contributions of these interactions to gut microbiota function in health and disease remain unclear.

To quantify this relative contribution, we introduce a *net interaction* metric, *ρ*, defined as the difference between the total cross-feeding and competition interactions within the community, normalized by the sum of all interaction strengths (see Eq. S8 in Methods (*86*)). Since this metric is normalized, the information it provides remains independent of community size and, consequently, diversity.

Our analysis reveals that the healthy state in the model is characterized by a negative net interaction (i.e. community dominated by competition), whereas the dysbiotic state is characterized by positive net interactions (cooperation-dominated community, Fig. 3A), which we use to rigorously identify the two states in the model (see Methods). This observation leads us to hypothesize that in IBD and potentially in other diseases, the gut microbiome may shift toward greater cooperation compared to the dynamics seen in healthy controls. While the net interaction metric robustly captures this transition, diversity metrics such as the Shannon index are broadly distributed for any given value of *ρ* (Supplementary Fig. S8), which may explain why it fails to consistently distinguish healthy from diseased states across conditions.

**Figure 3:**
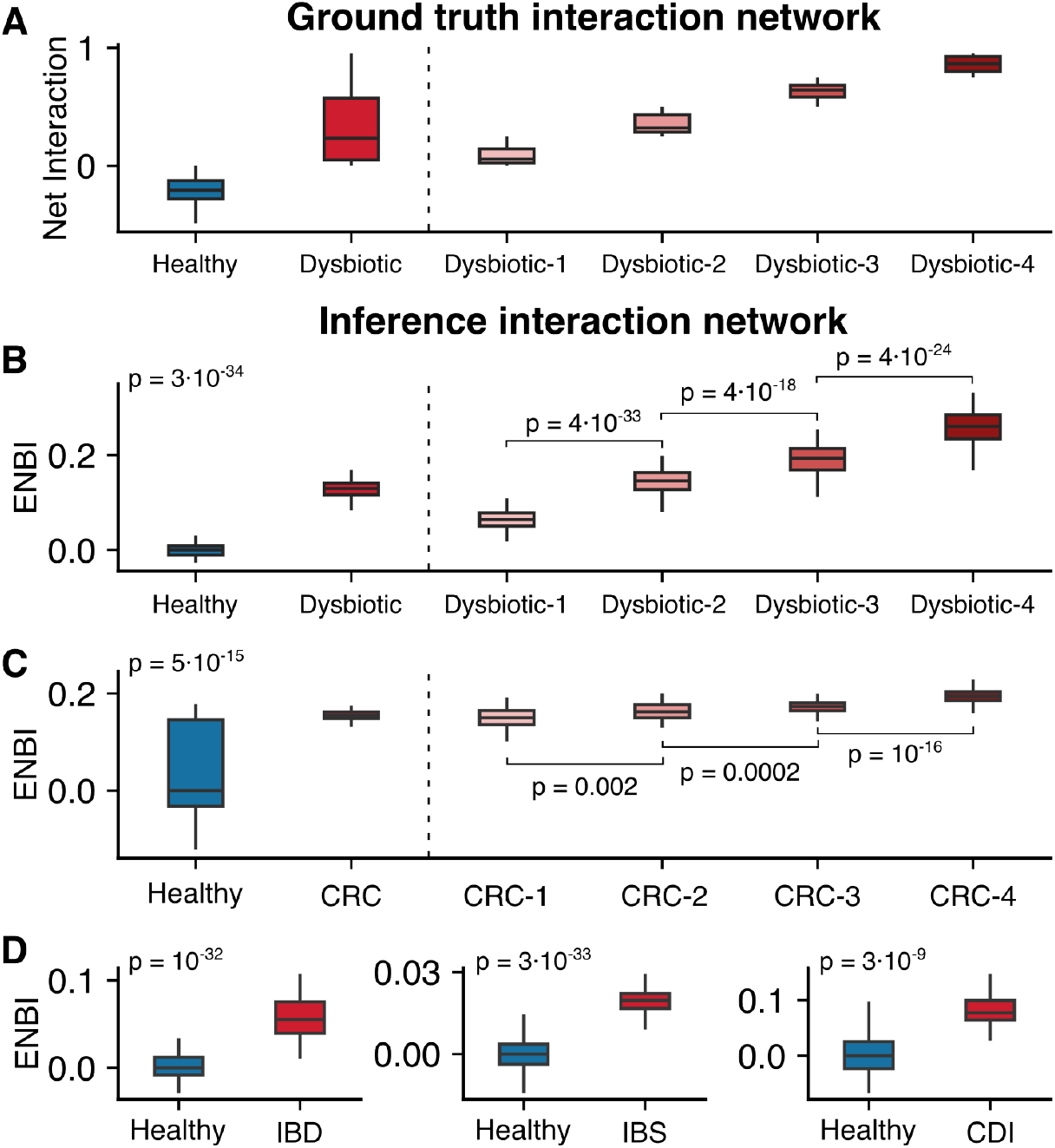
The Ecological Network Balance Index (ENBI) discerns healthy from pathological communities, and correlates with disease progression. (**A**) Net interaction calculated from the actual interaction network in the model, for healthy and dysbiotic states separately (left side of the dashed line) and different consecutive snapshots of unhealthy states (right side). For the latter, snapshots were grouped and ordered in 4 sets (i.e. quartiles) from smaller to larger *ρ*. (**B**) ENBI calculated for model data, showing that this indicator captures the same qualitative behavior as the net interaction. (**C**) ENBI for healthy and CRC patients (left side of dashed line) showing that the latter have a positive ENBI, which suggests a more cooperative microbial community compared to healthy patients; the ENBI increases with disease progression (right side), supporting our hypothesis of a progressive shift towards more cooperative network structures. (**D**) ENBI for IBD, IBS, and CDI also show that diseased states tend to be more cooperative than their respective controls.

To test our transition hypothesis, we used an algorithm for empirical co-abundance networks to infer effective interaction networks (see Methods (*86*) and (*104*)) and calculated the associated net interaction metric, *ρ*_inferred_, for available real-world cross-sectional data and our model-generated data. The latter allowed us to validate the inferred interaction network against the actual one.

As Fig. 3A and Supplementary Fig. S9A show, the net interaction *ρ*_inferred_ inferred for model snapshots successfully captured the qualitative behavior of the true underlying network, as *i*) simulated communities from dysbiotic states showed a higher *ρ*_inferred_ than those from healthy states; and *ii*) dysbiotic state communities with higher true-*ρ* values showed higher values of *ρ*_inferred_. Importantly, Fig. S9A shows that the inference method remains robust across different subsampling strategies. Building on this validation, we then applied this methodology across different diseases (IBD, CDI, IBS, and CRC). Supplementary Fig. S9B-E shows that, in all these cases, there is a significant increase in *ρ*_inferred_ compared to their respective healthy control groups. The qualitative results remained robust across additional network inference methods, one designed for cross-sectional data (*105*) (Supplementary Fig. S10) and another one, specifically developed for longitudinal data (*106*) (Supplementary Fig. S11). The agreement reinforces the association of dysbiosis with a shift toward a more cooperative microbial network.

With this in mind, we used the healthy/control case as a baseline for each dataset and computed the difference between the *ρ*_inferred_ values of the dysbiotic and control inferred networks, which we termed the “*Ecological Network Balance Index*” (ENBI) Eq. (S10). This index is negative when the community exhibits more competitive dynamics than the control (ENBI < 0) and positive when the community shows more cooperative interactions than in the control (ENBI > 0).

Across all diseases analyzed, as well as in our model, the dysbiotic state consistently exhibits a significantly positive ENBI value, reflecting a shift toward increased cooperative interactions compared to the controls (Fig. 3B-D). Furthermore, in the case of CRC, for which long-term data on disease progression is available, the ENBI increases (and thus community dynamics become increasingly cooperative compared to those in healthy individuals) as the disease advances (Supplementary Fig. S9B and Fig.3D). This consistent rise in the ENBI mirrors that observed in the model as the simulated community becomes more and more dominated by the few strains that compose the dysbiotic state.

## 5 Discussion

Whether substantial changes in the gut microbiome cause or result from the onset of disease remains debated. Regardless of causality, our study sheds light on the ecological dynamics and community-level effects shaping the gut microbiome in health and disease. The network ecological balance index (ENBI), our newly introduced metric to quantify the dominant interaction dynamics, offers deeper and more nuanced insights into the structure of microbial communities and underlying mechanisms than existing biomarkers. Unlike traditional metrics (e.g. Shannon index, which is abundance-based), the ENBI consistently reveals a reproducible pattern across diseases and methodologies, highlighting a microbial shift toward more cooperative dynamics in dysbiotic states compared to healthy controls.

In particular, our model suggests that diseased states are associated with a shift in microbial interaction networks, by which cooperative dynamics increasingly dominate over the competitive interactions. This relative increase does not necessarily reflect an absolute rise in cross-feeding; it could instead result from a sharper decline in competitive interactions. Thus our observation is not in conflict with previous reports of reduced cross-feeding and increased metabolic self-sufficiency in dysbiosis (*102,103, 107*). Furthermore, prior studies typically focus on specific metabolic pathways that get altered in disease, whereas our analysis assesses the overall balance between interaction types.

A detailed analysis of the resulting microbial communities indicates that this shift stems from the formation of mutualistic consortia, typically comprising a subset of species from the more heterogeneous healthy state. These consortia are more efficient at exploiting available resources and thus outcompete other species, potentially depleting metabolites, all of which may lead to the alteration or even loss of functions essential for host health. This exclusion often, but not always, results in reduced *α*-diversity, which explains why some diseases do not lead to a reduced diversity with respect to the control case.

Moreover, for colorectal cancer —for which data on disease progression are available— the ENBI correlates with disease advancement. If this trend holds across other diseases, the finding carries significant clinical implications, particularly because our metric can be quantified non-invasively from stool metagenomic data. Therefore, monitoring the ENBI of a patient can help devise early-warning indicators for disease onset, especially important for conditions such as CRC for which early detection is key to the success of the treatment and improvement of survival rates (*108*). A potential detection protocol could involve an initial ENBI calibration through regular stool samples over a set period to establish a patient-specific baseline interaction network, as part of annual health checkups. Subsequent, less-frequent sampling could then track significant ENBI deviations, signaling the emergence of dysbiosis. Even if symptoms are already present when monitoring starts, the rate of ENBI change would indicate disease progression. Nonetheless, despite the broad applicability of the ENBI, the biomarker by itself does not identify the specific disease causing dysbiosis. Thus, to maximize its diagnostic potential, the ENBI must be combined with other diagnostic tools for precise disease identification (e.g., for CRC, computed tomography colonography, endoscopy, or specific biochemical markers).

Beyond its diagnostic potential, this framework offers a tool to explore how factors such as diet, probiotics, lifestyle, antibiotics, or fecal microbiota transplantation influence the ecological interactions that shape the microbiome. In doing so, it opens up intriguing questions and research directions: Is the success of FMT in conditions like CDI linked to its ability to restore competition-driven microbial dynamics? Do probiotics work by re-establishing competitive interactions, or by preventing the emergence of dominant pathological consortia? How do diet and lifestyle contribute to shaping microbial communities? Can insights into ecological interaction networks obtained with our framework be leveraged to develop targeted interventions aiming to restore dysbiotic communities to a healthy state?

Our model follows a metabolically explicit approach but, unlike genome-scale metabolic models and kinetic models (*64*), which require detailed genome annotations and well-defined objective functions with extensive parameter fitting, our simpler framework focuses on essential ecological and metabolic principles and a biologically plausible parametrization. Our bottom-up approach is thus particularly well-suited for unveiling mechanistic underpinnings of gut microbiome emergent properties such as diversity, multistability, and metabolic organization (*65*). For example, our model provides an explanation for the Anna Karenina Principle (AKP, the observation that the microbiomes of healthy individuals are seemingly similar to each other, but dysbiotic communities are all distinct from each other; Supplementary Fig. S6): on the one hand, healthy states show similar functional composition, even though the communities able to produce those functions may be different; on the other hand, the loss of metabolic pathways disrupts partially or totally some of those functions in dysbiotic states, with disease data suggesting that the disruption may be disease-specific (*12*). Moreover, the model suggests that, for the loss of a function, not only the identity of the missing bacterial types matter but also the consequential changes in network structure. The subtle behavior for *β*-diversity observed in the model, and lack of consistency observed in our analysis of the data, agree with the lack of generality of the AKP across diseases (*109*).

Nonetheless, our framework is not without limitations. The model includes simplifications such as the indirect (as opposed to explicit) representation of anabolic reactions; the representation of host influence solely through resource input and dilution; and the reduced parametrization for energy and enzymatic cost matrices. These are in addition to other factors traditionally overlooked in models such as the dynamic interaction with phages or chemical warfare between bacteria. Expanding the model to incorporate a more realistic representation of these aspects is an important avenue for future work.

Furthermore, our approach to analyzing empirical data relies on inferred ecological interactions, which intrinsically depend on data quality and quantity and come with inherent assumptions and challenges (*63, 110, 111*). Other limitations are inherent to the heterogeneity of the methods used across studies. For example, cross-disease comparisons —which could provide valuable insights into microbiome dynamics for different conditions— are currently hindered by variability in sampling and taxonomic profiling methods across studies. Such methodological differences directly impact the inference of the ecological interaction network, making quantitative comparisons across diseases difficult. For that reason, we focused here on identifying qualitative trends rather than absolute numerical differences. Establishing standarized unified sampling and profiling methods would enable more robust cross-disease analyses in the future.

Addressing the challenges above will require a multidisciplinary effort that integrates empirical studies with theoretical approaches like the one we presented here. Connecting microbial ecology with clinical research, our framework contributes to precision medicine. By uncovering links between ecological interactions, disease progression, and therapeutic outcomes, we hope to inspire further research and more personalized strategies for maintaining or restoring gut health.

## Acknowledgments

The authors would like to thank Jacopo Grilli, Matteo Sireci, and the members of the Bonachela lab for helpful discussions.

## Funding

We acknowledge the Spanish Ministry of Research and Innovation and Agencia Estatal de Investigación (AEI), MICIN/AEI/10.13039/501100011033, for financial support through Project No. PID2023-149174NB-I00, funded also by ERDF/EU, and Project No. PID2020-113681GB-I00 funded by MICIN/AEI/10.13039/501100011033. This work was also supported by a Simons Early Career Investigator in Marine Microbial Ecology and Evolution Award (award #826106) and NSF grant DMS-2052616 to J.A.B.

## Author contributions

JAB and MAM designed and supervised the research. RCL, JAB, and MAM developed the model, with input of SAL, MJB, MM, and MGDB. RCL collected the data and performed statistical and modeling analyses. All authors wrote and reviewed the manuscript.

## Competing interests

There are no competing interests to declare.

## Data and materials availability

No empirical data were produced in this study. All the publicly available datasets are cited in the section “Datasets” in (*86*). Our code will be available on either zenodo or github.

## Supplementary Materials

## Materials and Methods

### Measures

#### p-value

We assessed statistical significance using the Mann-Whitney U test (two-sided).

#### *α*-Diversity: measured using the Shannon Index, defined as

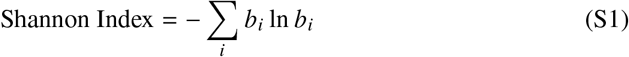

where *b*_*i*_ represents the relative abundance (biomass) of type *i* in the model, or the relative abundance (proportion of counts) of species *i* in the metagenomic data.

#### *β*-diversity

measured using the Bray-Curtis (BC) dissimilarity index between two samples, *a* and *d*, defined as:

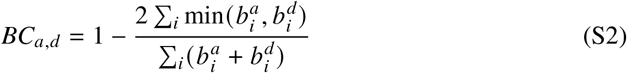

where 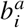 and 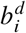 represent, for samples *a* and *d*, the relative abundance of type *i* (model) species *i* (metagenomic data). A value of *BC*_*a,d*_ = 0 indicates identical community compositions, while *BC*_*a,d*_ = 1 indicates completely distinct communities.

#### Number of Enzymes

In real data, we calculated the number of enzymes as the sum of enzymes with non-zero abundance in the sample. In the model, because enzymes are present through the enzymatic cost of a pathway, we calculated a proxy for the number of enzymes, the total enzymatic cost in the sample:

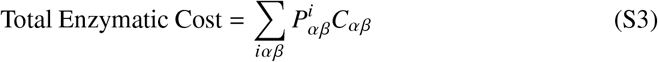

where 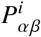 represents the pathway matrix of bacterial type *i* and *C*_*αβ*_ the associated cost stemming from enzyme production.

#### Number of Metabolites

Calculated as the sum of non-zero metabolites

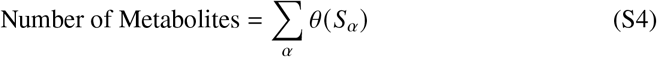

where *θ* (*x*) is the Heaviside function (i.e. Θ(*x*) = 1 if *x* > 0, and Θ(*x*) = 0 otherwise) and *S*_*α*_ is the density of metabolite *α*, both in the model and metabolomic data.

#### Pathway fraction

In real data, the pathway fraction is computed as the relative abundance of each metabolic pathway. In the model, we define it as:

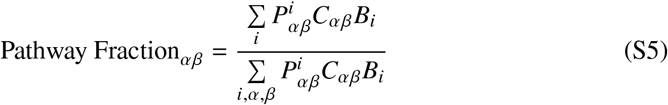

This formulation incorporates both the enzymatic cost and the biomass of the species expressing each pathway, providing a proxy for the functional signal that would be obtained from pathway abundance profiles in metagenomic data.

#### Cross-feeding

calculated as

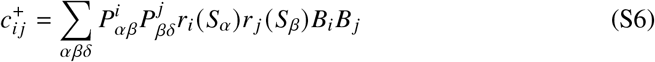

which captures all the cross-feeding interactions from type *i* to type *j*, mediated by all possible metabolic pathways. Specifically, 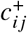 considers all instances where species *i* consumes substance *S*_*α*_ to generate substance *S*_*β*_, which is then consumed by species *j* to produce any byproduct *S*_*δ*_. Each interaction is weighted by the corresponding species abundances (*B*_*i*_ and *B* _*j*_) and consumption factor *r*_*i*_, the latter given by the Monod function 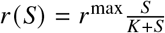. The matrix composed of all 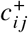 is always positive and non-symmetric.

#### Competition

calculated as

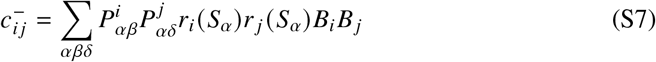

which captures the competition between types *i* and *j* arising from their shared preference for the same substance *S*_*α*_. Each competitive interaction is weighted by the corresponding species abundances and consumption functions. The matrix composed using all 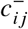 is always positive and symmetric.

#### Net interaction

defined as

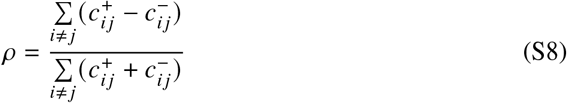

where the terms 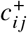 and 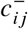 represent, respectively, the elements of the cross-feeding and competition matrices above.

#### Inferred net interaction

defined as

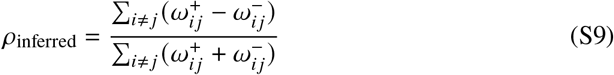

where *ω*_*i j*_ represent the positive 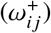 or negative 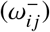 weights between types (in the model) or species (metagenomic data) *i* and *j* in the inferred interaction network. See section “Inferring the interaction network” below for details.

#### Ecological Network Balance Index (ENBI)

caculated as

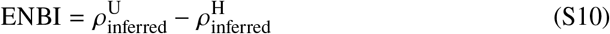

where *ρ*_inferred_ are the unhealthy (dysbiotic, 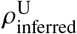) and healthy (healthy, 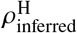) inferred net interactions for the different cases analyzed (diseases and model).

### Definition of healthy and dysbiotic states

To classify states beyond a simple visual inspection of the temporal dynamics (e.g. Fig. 2B), we define the **healthy state** as the configuration of the system in which the net interaction among present bacteria is negative (*ρ* < 0). Conversely, the **dysbiotic state** is characterized by a positive net interaction among bacteria in the system (*ρ* > 0).

We examined multiple alternative definitions based on other biomarkers, including combinations of the Shannon index, richness, and the number of substances or metabolites, among others. However, the criterion above proved to be the definition most in agreement with the classification of states obtained through visual inspection.

### Inferring the interaction network

To infer the underlying interaction network of the microbial community, we used three complementary methods: two suitable for cross-sectional datasets (i.e., data from different subjects) and another one specifically devised for longitudinal datasets (i.e., data collected over multiple time points from the same subject).

#### Method 1. *FlashWeave*

This algorithm infers interaction networks from cross-sectional data using co-abundance networks. These networks are built on the principle that bacterial species with positively correlated abundances likely exhibit positive interactions, while negatively correlated abundances suggest negative interactions. Importantly, the algorithm goes beyond simple correlation by incorporating ways to filter out indirect interactions mediated by other species, and distill direct ones. See (*104*) for further details.

As output, this algorithm generates a single undirected interaction network for a given set of samples. To obtain statistical insights, we used bootstrapping (or subsampling) on each dataset. The latter involves generating multiple subsamples from the full dataset and inferring the interaction network for each subsample. For instance, Fig. S9 shows the inferred net interaction, *ρ*_inferred_, for 500 bootstrapped subsamples across all datasets, including both model outputs and various disease datasets, using fixed fractions of the total sample set. Additionally, Fig. 3 displays the results for a specific subset obtained with a fraction of 0.8. Across all cases, we consistently observed that the net interaction of the inferred network is higher in diseased (dysbiotic) states compared to control (healthy) states for both real datasets and the model, respectively.

For the datasets corresponding to different diseases (IBD, IBS, CDI, CRC), we used the *sensitive* and *non-heterogeneous* options of FlashWeave in conjunction with its default parameters (*104*). For the data generated with the model, where the number of samples exceeds one thousand, we applied the *heterogeneous* option, recommended for large datasets in the original publication.

#### Method 2. *BEEM-Static*

This algorithm, which we used as an additional validation of the results obtained with method 1, infers ecological interactions from cross-sectional data by estimating the parameters of a generalized Lotka-Volterra (GLV) model. *BEEM-Static* explicitly models species interactions by assuming that microbiome communities are at or near equilibrium. Specifically, the method employs an expectation-maximization algorithm to iteratively estimate both species’ biomasses and interaction strengths, filtering out samples that violate the equilibrium assumptions. Notably, this approach enables the inference of directed interaction networks and has been shown to outperform standard correlation-based methods in capturing both the presence and directionality of microbial interactions.

The algorithm includes various tweaking parameters, which are detailed in the original publication (*105*). In our implementation, we adhered to the default settings with two exceptions: we increased the maximum number of iterations within the model to 50 and applied preprocessing steps to our datasets. Specifically, we used from the datasets only bacterial taxa that were present in at least 30% of the samples and had a mean relative abundance greater than 10^−4^, as the algorithm did not seem to work otherwise. As with Method 1, we used bootstrapping to assess statistical significance. Supplementary Fig.S10 presents the results of our subsampling approach. Consistent with the findings from Method 1, the figure shows that the net interaction of the inferred network is systematically higher in diseased (dysbiotic) states compared to control (healthy) states in both real datasets and model-generated data.

#### Method 3. *LIMITS*

This algorithm leverages a discrete-time Lotka-Volterra model to infer undirected interaction networks from longitudinal data. The algorithm includes three key tweaking parameters: the number of species considered for network inference and two parameters specific to the algorithm (the error threshold and the bagging –or bootstrap aggregation– determining the number of internal iterations performed). While the inference process is relatively straightforward, we refer the reader to the original publication for a detailed explanation (*106*).

For each subject, we selected a predefined number of species and infer the interaction network across various error thresholds from 1 to 5 (ranges employed in the original publication). We used the algorithm over a large number of iterations (10^6^) (real data) and 5 · 10^4^ (model data) to ensure robustness, although we noticed that the iteration count did not significantly affect the results. The final network was constructed by first averaging the outcomes from all iterations corresponding to each error threshold, and subsequently averaging the results across all error thresholds. In our implementation, we applied a slight modification to the original algorithm by using the average, rather than the median, of all iterations to construct the final network. This modification allowed us to obtain more consistent results.

### Model simulation details

We integrated numerically the model equations described in the Supplementary Text by using a multi-step variable-order integration scheme available in Python (*solve_ivp*), in which we adapted the integration time steps to prevent negative abundances. The parameters used for all simulations (unless otherwise specified) can be found in Table S1. The enzymatic cost matrix, *C*_*αβ*_, is assumed to be identical to the energy matrix, *E*_*αβ*_, as differences are inherently accounted for through the non-linear trade-off function (see Supplementary Text Eqs. S14 and S15). The trade-off function used for the simulations presented in the main text used parameters *b* = 8, *c* = 6, and *v* = 1.1.

### Functional coarse-graining

In the model, coarse-graining is implemented by clustering substances with similar energy levels into functional groups (see Fig. S2 for illustration). Let us consider the set of all substances *S*_*α*_ which, as explained in Supplementary Text, is ordered by energy level. For fine coarse-graining, substances are grouped in sets of four, meaning that *S*_1_-*S*_4_ are treated as a single functional entity, *S*_5_-*S*_9_ as another, and so on. An exception is made for *S*_0_, which is always considered separately since it is the only substance directly introduced into the system. Once substances are grouped, we consider all possible metabolic routes between these grouped entities, resulting in *n*_fg_ = 43 functional groups. For broad coarse-graining, a larger grouping size of 10 is used, leading to a more aggregated representation of metabolic pathways. In this case, the number of functional groups is reduced to *n*_fg_ = 9.

In the IBD dataset, we annotated and grouped metabolic pathways according to the BioCyc ontology classification (*112*). We assigned a different color to each functional group, with individual pathways represented by different shades within their respective groups. In Fig. 2G, these groups are ordered from bottom to top as follows: Carbohydrate Degradation (blue), Amino Acid Biosynthesis (orange), Nucleoside and Nucleotide Degradation (light green), Carbohydrate Biosynthesis (red), All Other Biosynthetic Pathways (yellow), All Other Pathways (purple), Generation of Precursor Metabolites and Energy (dark green), Cofactor, Carrier, and Vitamin Biosynthesis (brown), Fatty Acid and Lipid Biosynthesis (pink), Nucleoside and Nucleotide Biosynthesis (light blue), Cell Structure Biosynthesis (black).

### Datasets

For the analysis of diseases, we used the following datasets (data used as obtained from original sources):

#### IBD

Metagenomic, enzymatic and metabolomic data were obtained from the Inflammatory Bowel Disease Multi’omics Database (https://ibdmdb.org/).

#### IBS

Metagenomic and enzymatic data were obtained from (*44*). Metabolomic data obtained from (*113*).

#### CDI

Metagenomic data obtained from (*43*).

#### CRC

Metagenomic, enzymatic, and metabolomic data were obtained from (*48*).

#### F4 and M3 gut data

Metagenomic data obtained from https://github.com/twbattaglia/MicrobeDS.

#### Model data

Temporal analyses and macroecological pattern assessments were based on weekly sampled data from our simulations. For analyses comparing healthy and dysbiotic states, we used 1000 samples per state (or the maximum number of samples available) for each realization, with approximately 100 independent realizations included in the analysis.

**Table S1:** Experimental values of the model parameters. Model parameter values.

| Symbol | Definition | Value | References |
| --- | --- | --- | --- |
| $\gamma$ | Energy to biomass conversion | $1.05 \times 10^{13}$ cell/mol ATP | (114, 115) |
| $\delta$ | Outflow rate | $\frac{1}{24}$ 1/h | (116) |
| $h$ | Inflow rate of substances | 2 g/(L h) | (72, 117, 118) |
| $K$ | Half-saturation constant | $10^{-4}$ g/L | (119, 120) |
| $r^{\max}$ | Maximum uptake rate | $10^{-10}$ g/(L cell h) | (120) |
| $U$ | Invasion rate | $\frac{2}{3}$ invasions $\text{h}^{-1}$ | - |
| $e_{\max}$ | Maximum energy per resource consumption | 3 mol ATP/mol of S | - |
| $M$ | Molar mass of substances | 100 g/mol | - |
| $V_{\text{colon}}$ | Volume of the colon | 0.41 L | (121) |
| $R$ | Total number of substances | 30 | - |
| $B_T$ | Total biomass in a typical gut microbiome | $\sim 3.9 \times 10^{13}$ cell | (121) |

## Supplementary Text

### Model building

We model the gut microbiome as living in a chemostat-like environment, a common simplifying assumption (in reality, food intake and substance excretion occurs in a discrete, rather than continuous, manner). We model the ecological dynamics of the system using consumer-resource equations, also considering metabolic preferences. Specifically, we consider a number *N* (*t*) of bacterial types (e.g., species, strains, or another suitable taxonomical group) with abundances *B*_*i*_ (*t*), where *i* identifies the bacterial type. Additionally, we include a maximum number of possible resources (or substances, as we use indistinguishably the two terms here) *R*, with concentrations *S*_*α*_, where *α* indicates the type of substance. Substances are introduced into the system at an inflow rate *h*_*α*_ and may also be generated as byproducts of bacterial metabolism. New bacterial types are introduced at a certain rate *U* with a small initial relative abundance of 10^−4^, as long as their invasion fitness —initial per-capita growth rate of the invader in the environment created by the existing bacterial types— is positive. If the invasion fitness is negative and the initial biomass of the invading bacteria is sufficiently small, the new bacterial type is declared extinct in the next temporal step without affecting the system. Therefore, only viable invader types contribute to the dynamics of the system.

We explicitly incorporate metabolic pathways into the model by representing all possible routes between any two distinct substances. To describe the metabolic pathway that converts a substance *S*_*α*_ into another substance *S*_*β*_, we introduce two key quantities:

- The energy *E*_*αβ*_ = *E*_*α*_ − *E*_*β*_, or benefit, derived from the route, where *E*_*α*_ and *E*_*β*_ are the energies associated with substances *α* and *β*, respectively. This *E*_*αβ*_ reflects the free energy released during the reaction, energy that we assume is fully available for growth. Nonetheless, this assumption could be relaxed by incorporating a conversion constant to account for partial energy utilization.
- The enzymatic cost, *C*_*αβ*_, required to complete the pathway, which accounts for the number and complexity of the enzymes involved.

For simplicity, we explicitly focus on catabolic reactions, i.e. those in which the free energy of the reaction is positive, and account for anabolic processes indirectly only (see below). In particular, we assume that all metabolic routes where *E*_*αβ*_ > 0 (i.e. catabolism) are possible, and that the energy obtained and the enzymatic cost needed for the route are the same (see below for details on how this is implemented in the model via the trade-offs). Nonetheless, the framework is versatile and can accommodate more complex and realistic metabolic structures through parameter adjustments.

Each bacterial species is characterized by its own specific metabolic pathways, represented by a binary matrix *P*^*i*^. In this matrix, 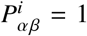 indicates that bacterium *i* can metabolize *S*_*α*_ into *S*_*β*_, while 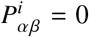 indicates it cannot. These pathways are randomly assigned when the bacterium is introduced into the system and remain fixed, therefore assuming no evolutionary adaptation occurs. First,a random number of pathways is assigned to the bacterium, ranging between 1 and a specified maximum. The maximum is chosen to be relatively small to reduce computational overhead, but sufficiently large to not limit the introduction of viable bacteria into the system. Then, the specific pathways are selected randomly from all possible *S*_*α*_ → *S*_*β*_ combinations, adhering to the catabolic constraint *E*_*αβ*_ > 0. This approach ensures a diverse yet computationally manageable set of pathways for each bacterium, reflecting ecological variability while maintaining system feasibility.

### Model equations

The equations representing the model are as follows:

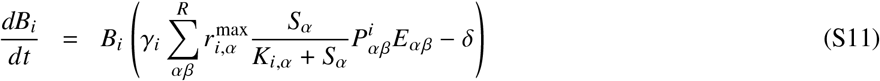

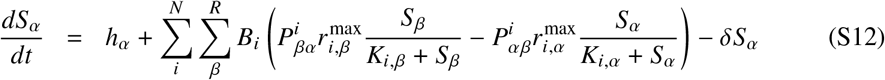

where *γ*_*i*_ is the yield or conversion constant from energy to biomass for bacterium *i*; 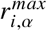 and *K*_*i,α*_ are, respectively, the uptake rate and half-saturation constant for bacterium *i* feeding on substance *α*; *δ* is the outflow rate; and *h*_*α*_ is the inflow rate for substance *α*. The values for all parameters are fully described in Table S1. We assume growth parameters, *r*^max^ and *K*, to be the same across all bacterial types and substances, and that only the most energetic substance has a non-zero inflow rate. For convenience, we express the (unique) energy associated with a substance *α* with different units, using the conversion 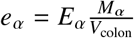; we further define such energy as a fraction of a maximum energy, i.e. *e*_*α*_ = *e*_max_(1 − *α*/*R*).

Finally, to ensure a balanced and realistic pathway structure within the system, a trade-off is introduced, imposing disadvantages on bacteria with numerous and costly pathways. This trade-off prevents unrealistic scenarios in which a single bacterial type that shows all pathways ultimately dominates the community. The trade-off is implemented as a growth penalty proportional to the number and cost of pathways utilized by the bacteria:

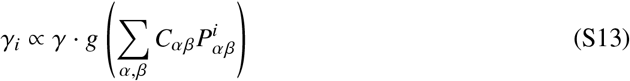

where *g* is a decreasing function. For the latter, we have explore the following two families of functions:

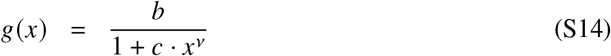

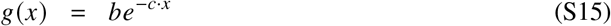

where *b, c* and *v* are three positive constants that help us vary the different trade-offs inside the same family, and verify the robustness of the main results.

### Influence of the trade-off

As shown in Fig. S3, the qualitative results of our model remain robust across different trade-off functional forms and parameterizations. Nonetheless, as expected given the nature of the tradeoff (see above), certain parametrizations can lead to ill-defined systems in which either no bacteria survive or only a single species dominates.

The trade-off primarily determines the total enzymatic cost allocated by each bacterial type and how this cost is distributed—either across multiple low-cost pathways or concentrated in fewer high-cost ones. Fig. S3F shows the distribution of total enzymatic cost per type across different trade-offs. We can approximate the peak of this distribution and gain insight into its width by considering the maximization of the per capita growth rate. First, we incorporate the trade-off function *γ*_*i*_ in the definition of the growth rate (Eqs. S11 and S13):

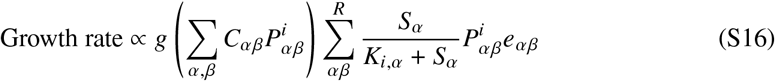

where 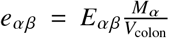 represents the energy of path *S*_*α*_ → *S*_*β*_ in different units (see above). Maximizing this equation exactly is not possible, because it depends on all substances in the system, which are in turn determined by the pathways of the existing bacterial community. However, assuming that each substance reaches approximately similar concentrations, we can simplify the expression to:

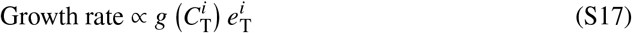

where 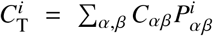 represents the total enzymatic cost per bacterial type, and 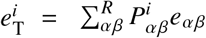 denotes the total energy a bacterial type *i* extracts from its pathways (up to a proportionality constant to maintain units). Given our assumption that the energy and enzymatic cost matrices are identical (or at least proportional), the optimal total enzymatic cost per type is given by the value that maximizes:

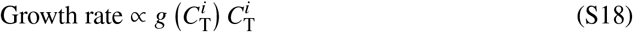

Fig. S3F shows, for different versions of the trade-off, the right hand side of this equation alongside the distribution of total enzymatic cost per type. As shown in the figure, the peak of this right hand side coincides with the peak of the distribution, whose width results from the fact that pathway configurations are assigned randomly and suboptimal pathways persist in the system for some time. This width, however, is also influenced by the trade-off as, when the right hand side of Eq.(S18) is flatter around its maximum, the resulting distribution is broader as suboptimal pathway configurations are similarly efficient to the optimal one, thus allowing for greater variability in enzymatic investment strategies.

In other words, if the maximum of Eq. S18 were significantly higher than the maximum energy obtainable per pathway, only bacteria with the most energetically efficient pathway would survive, leading to an ill-defined system. To obtain non-trivial communities, the trade-off function and parameters must ensure that the peak of Eq. S18 remains below the most energetic pathway. Moreover, the closer this maximum is to the mode of the distribution, the richer the microbial community will be, as a greater number of functionally distinct yet competitive enzymatic investment strategies can emerge.

### Simulation algorithm

We start our simulations with only one substance (the most energetic one, which is introduced in the system at a non-vanishing rate) and only one bacterial type able to metabolize this substance (and thus transform it into a random, different substance).

From that initial conditions, the simulations proceed according to the following algorithm:

0 Run the ecological dynamics, given by Eqs. S11-S12 until the next invasion step.
1. At each step, eliminate species that have fallen below the extinction threshold (set to be a fraction 10^−4^ of the total biomass, that is, an abundance 10^−4^ Σ_*i*_ *B*_*i*_).
2. Introduce an invader, a bacterial type whose pathways are randomly selected (see above).
3. Repeat until a selected ending time.

### Comparison to existing, relevant work

Recent studies have highlighted the importance of metabolic interactions in shaping microbial communities in health and disease. Marcelino et al. (*102*) introduced a quantitative framework for microbial cross-feeding, demonstrating that specific metabolite exchanges are disrupted in diseased states. Similarly, Watson et al. (*103*) and Veseli et al. (*107*) showed that microbes in dysbiotic conditions exhibit increased metabolic self-sufficiency, indicating reduced dependence on crossfeeding for certain anabolic pathways.

While these observations might initially appear contradictory to our results, they are in fact complementary. Our approach quantifies net metabolic interactions, incorporating both crossfeeding and competition. Unlike previous studies, which emphasize the loss of specific cross-feeding dependencies, our findings indicate that cross-feeding interactions become more significant relative to competition during dysbiosis. Importantly, this relative increase does not necessarily reflect an absolute rise in cross-feeding; it could instead result from a sharper decline in competitive interactions. Furthermore, prior studies typically focus on specific metabolic pathways that get altered in disease, whereas our analysis assesses the overall balance between interaction types.

These studies also emphasized how metabolic self-sufficiency provides an advantage under stressful conditions, aligning with our observation that disease states are characterized by fewer available metabolites, favoring taxa less reliant on external metabolic inputs. Marcelino et al. specifically suggest that high species diversity is associated with increased net metabolite production, creating more opportunities for consumer species to thrive. This aligns with our simulations, in which higher diversity consistently corresponds with greater metabolite availability.

Therefore, our results complement existing literature by illustrating that the balance of microbial interactions shifts during disease, notably elevating the relative importance of cross-feeding compared to competition. These findings highlight the value of integrative approaches that simultaneously consider both specific metabolic dependencies and the broader dynamics of microbial interactions.

**Figure S1:**
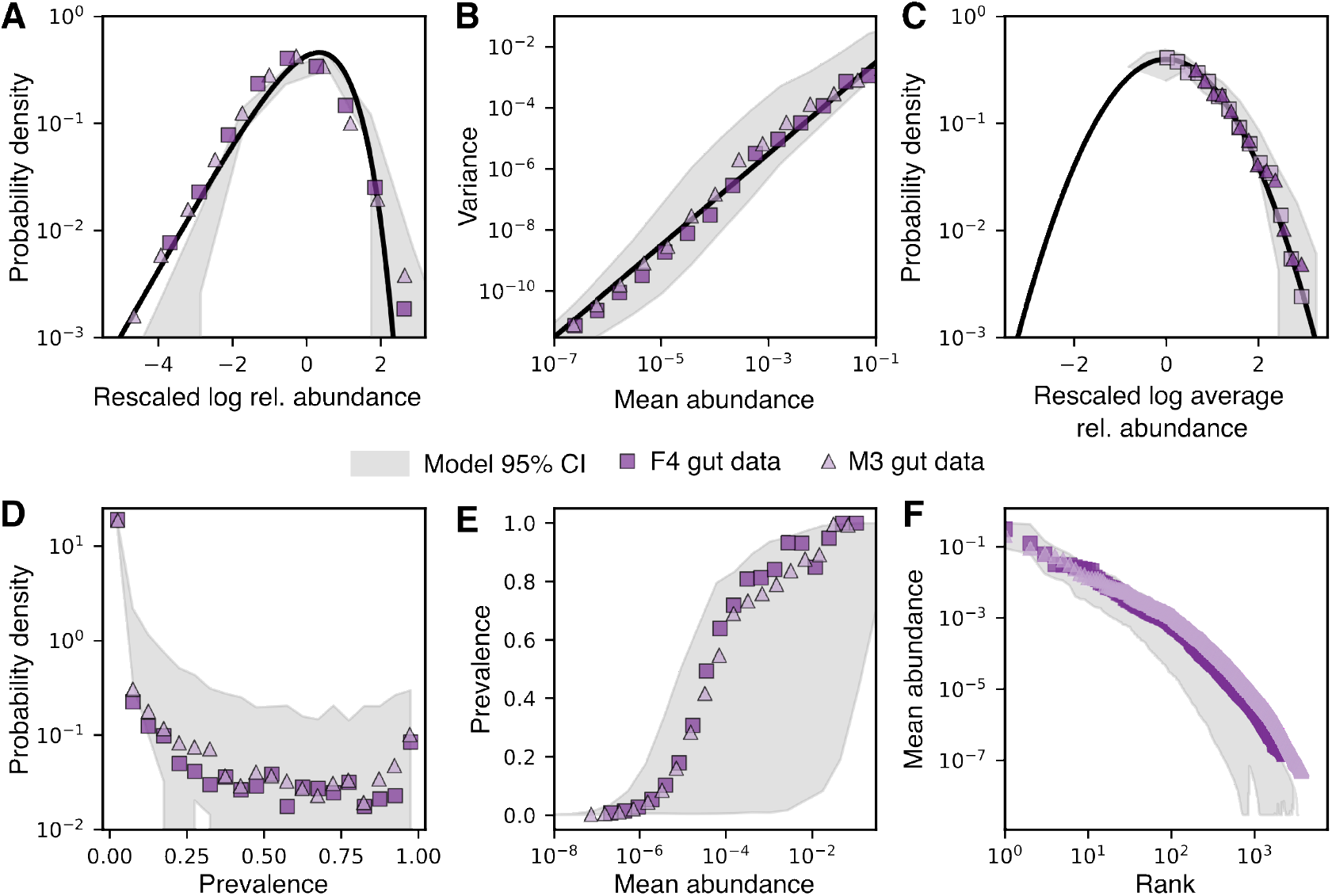
Model reproduces well-established macroecological patterns of the gut microbiome without any parameter fitting. **A-C**) Macroecological patterns described in (*80*). (**A**) The Abundance Fluctuation Distribution (AFD), which characterizes the distribution of a species’ abundances across samples. The solid black line represents the expected distribution obtained by rescaling the log of relative abundance (subtracting the mean and dividing by the standard deviation) when the relative abundance follows a gamma distribution. (**B**) The mean and variance of species abundance distributions follow Taylor’s Law, with variance scaling as a power of the mean, specifically with an exponent of 1.5 (black line). (**C**) The Mean Abundance Distribution (MAD), defined as the distribution of species’ mean abundance across samples, follows a lognormal distribution (black line). (**D-F**) Macroecological patterns described in (*81*). (**D**) Distribution of species prevalence, defined as the fraction of samples in which a species is present. (**E**) Relationship between species prevalence and mean abundance. (**F**) Rank distribution of mean abundances.

**Figure S2:**
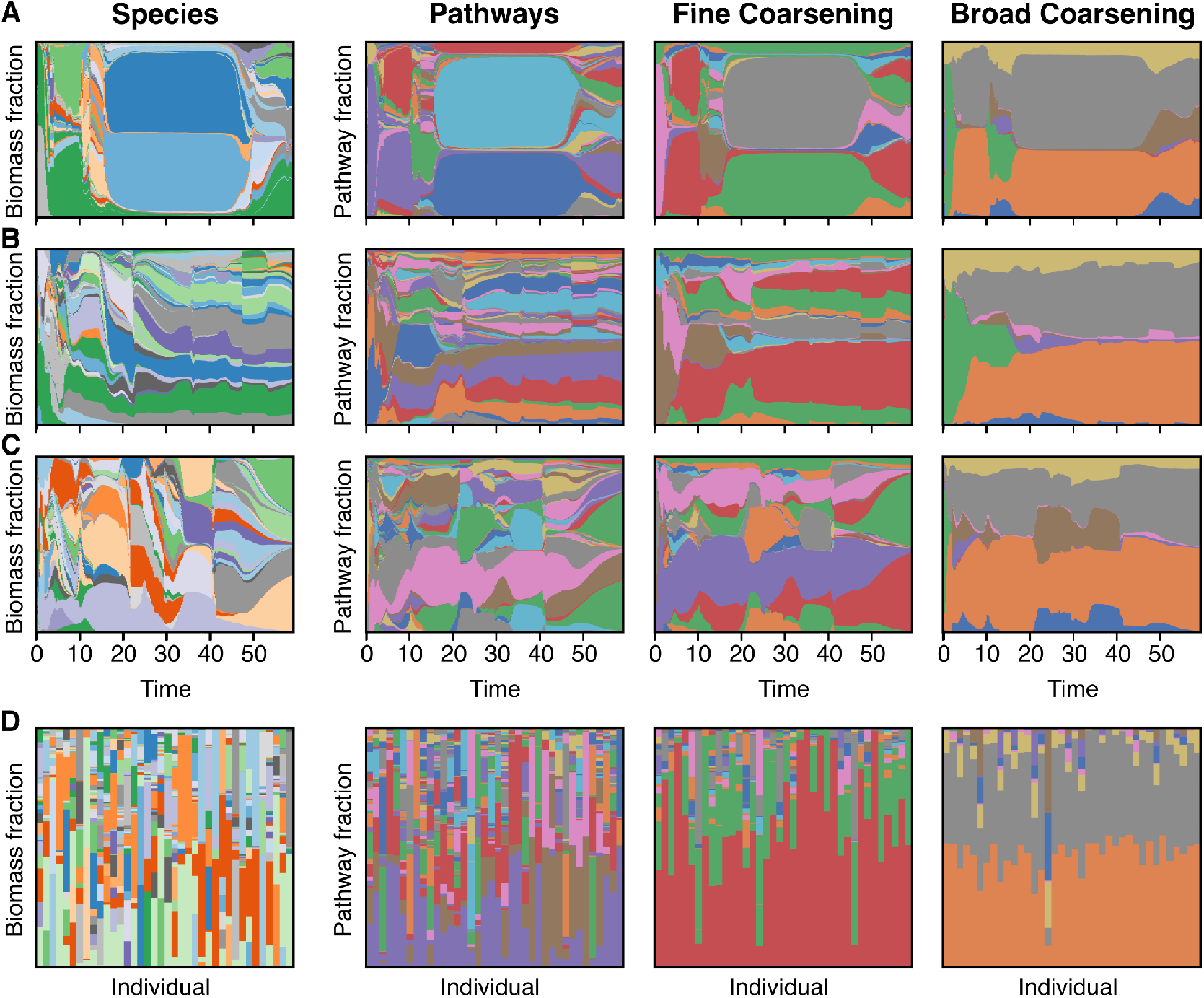
Model data reproduces functional redundancy. (**A-C**) Time-series of relative species abundance (left), enzymatic cost fraction for all pathways (number of resulting functional groups, *n*_fg_ = 435; middle left), for a fine coarsening (*n*_fg_ = 43; middle right), and for a broad coarsening (*n*_fg_ = 9; right) across three different realizations. (**D**) Transversal (across individuals) evaluation of the same metrics: biomass fraction at the species level and enzymatic cost fraction under different coarsening schemes. Coarse-grained pathways exhibit functional redundancy, meaning that taxonomically distinct species share similar functional profiles.

**Figure S3:**
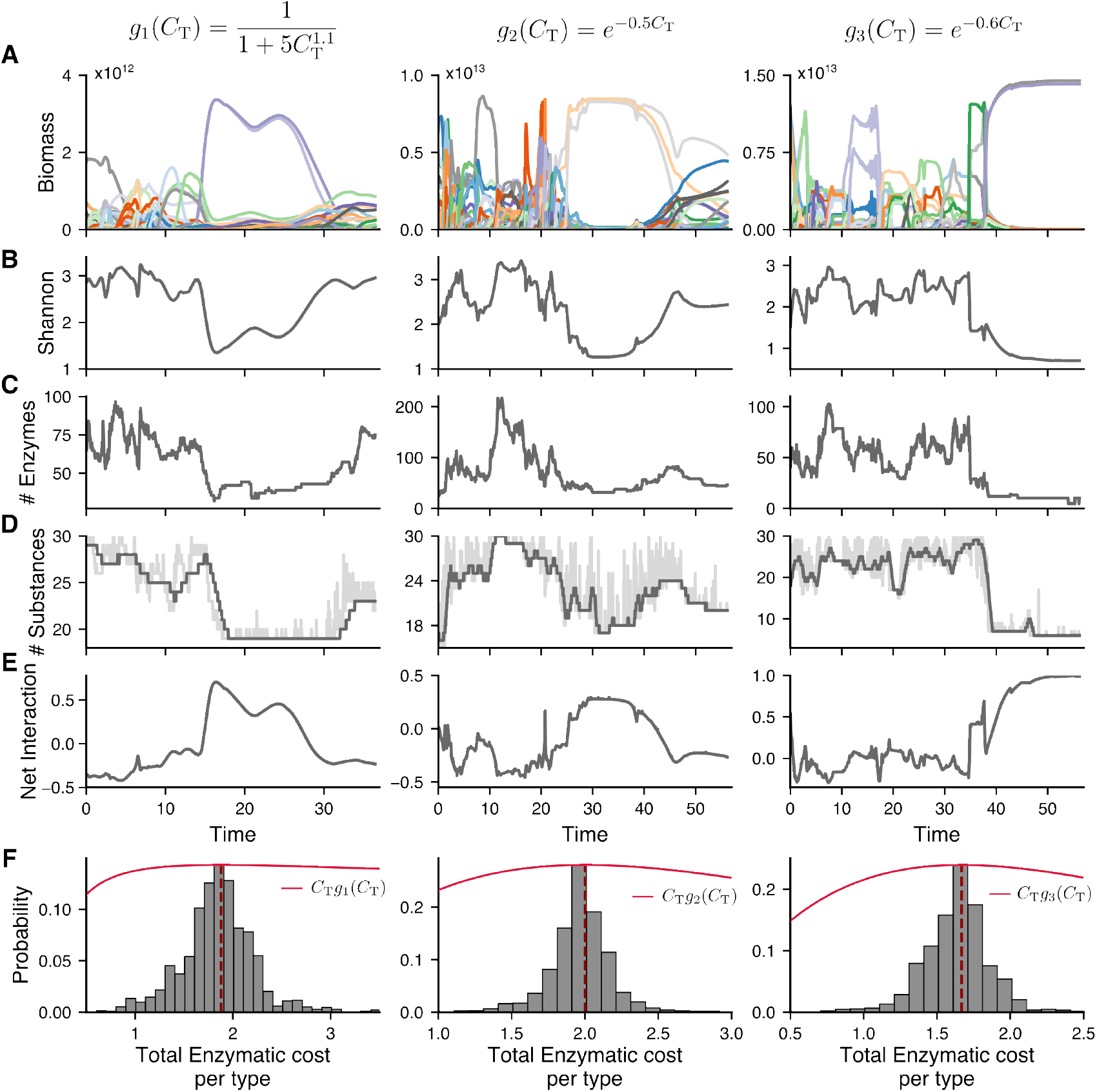
Alternative stable states are robust against trade-off changes. For three different trade-off functions, *g*_1_ (left), *g*_2_ (middle), and *g*_3_ (right), the following are shown: (**A-E**) Time-series of bacterial abundances (**A**), Shannon index (**B**), enzymatic cost (**C**), number of substances (**D**), and net interaction (**E**). (**F**) The total enzymatic cost per type (*C*_*T*_) distribution shows a peak at the maximum of *C*_T_*g*_*k*_ (*C*_T_), suggesting an optimal enzymatic investment strategy. The consistency of these patterns across different trade-off formulations and parameter choices indicates the robustness of the alternative stable states.

**Figure S4:**
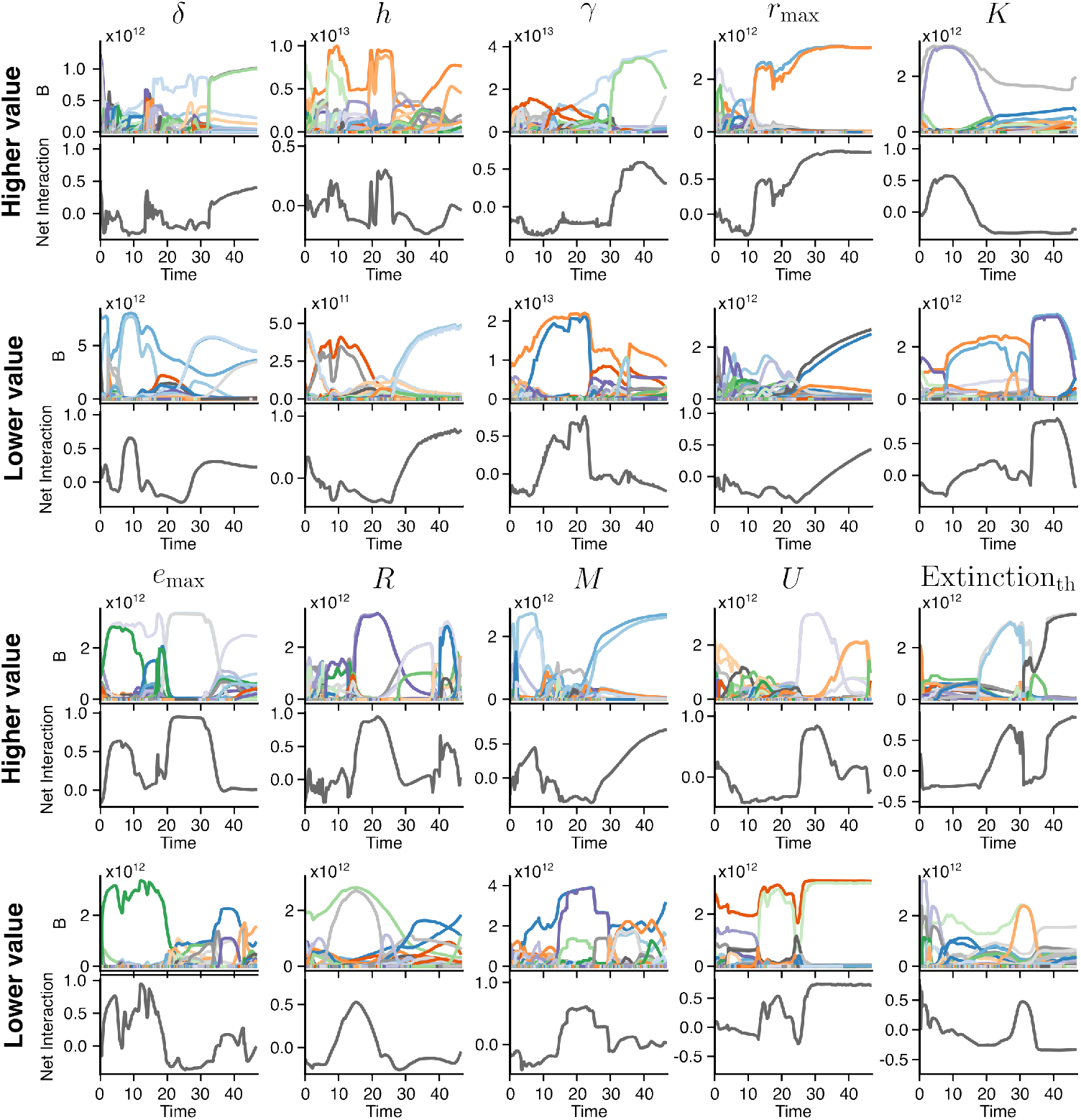
Alternative stable states are robust to parameter variations. Time-series of total biomass (top row in each subpanel) and net interaction (bottom row in each subpanel) for different parameter settings in the model. Each column represents a change from the original parameter values reported in Table S1. **(A)** Increased values: from left to right, *δ* = 2.5/24, *h* = 10, *γ* = 1.25 · 10^14^, *r*_max_ = 10^−9^, *K* = 10^−3^. **(B)** Decreased values: from left to right, *δ* = 0.4/24, *h* = 0.5, *γ* = 10^13^, *r*_max_ = 10^−11^, *K* = 10^−5^. **(C)** Increased values: from left to right, *e*_max_ = 4, *R* = 40, *M* = 120, *U* = 4, extinction threshold = 5 · 10^−4^. **(D)** Decreased values: from left to right, *e*_max_ = 2, *R* = 20, *M* = 80, *U* = 1, extinction threshold = 5 · 10^−5^. In all cases, the model continues to exhibit both healthy and dysbiotic states, demonstrating that the emergence of alternative stable states is qualitatively robust to wide parameter changes.

**Figure S5:**
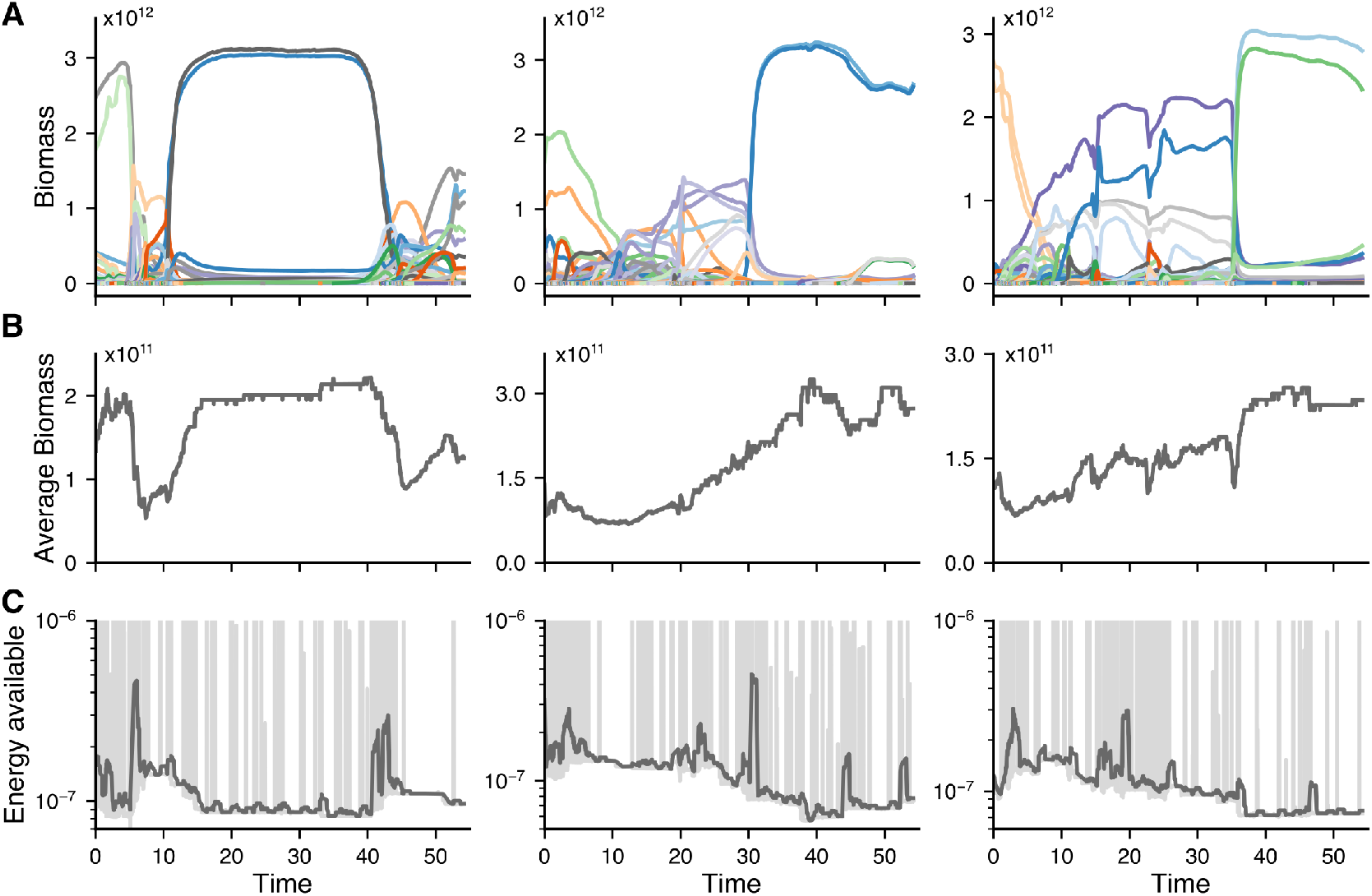
dysbiotic state shows greater efficiency than the healthy state. (**A-C**) Time-series of bacterial biomasses (**A**), average biomass across the system (**B**), and available energy within the system (**C**) for three different realizations of the model. In (**C**), dark grey lines represent the rolling average, while light grey lines depict the actual values. dysbiotic states show higher average biomass while depleting more available energy in the system, indicating a more efficient microbial community.

**Figure S6:**
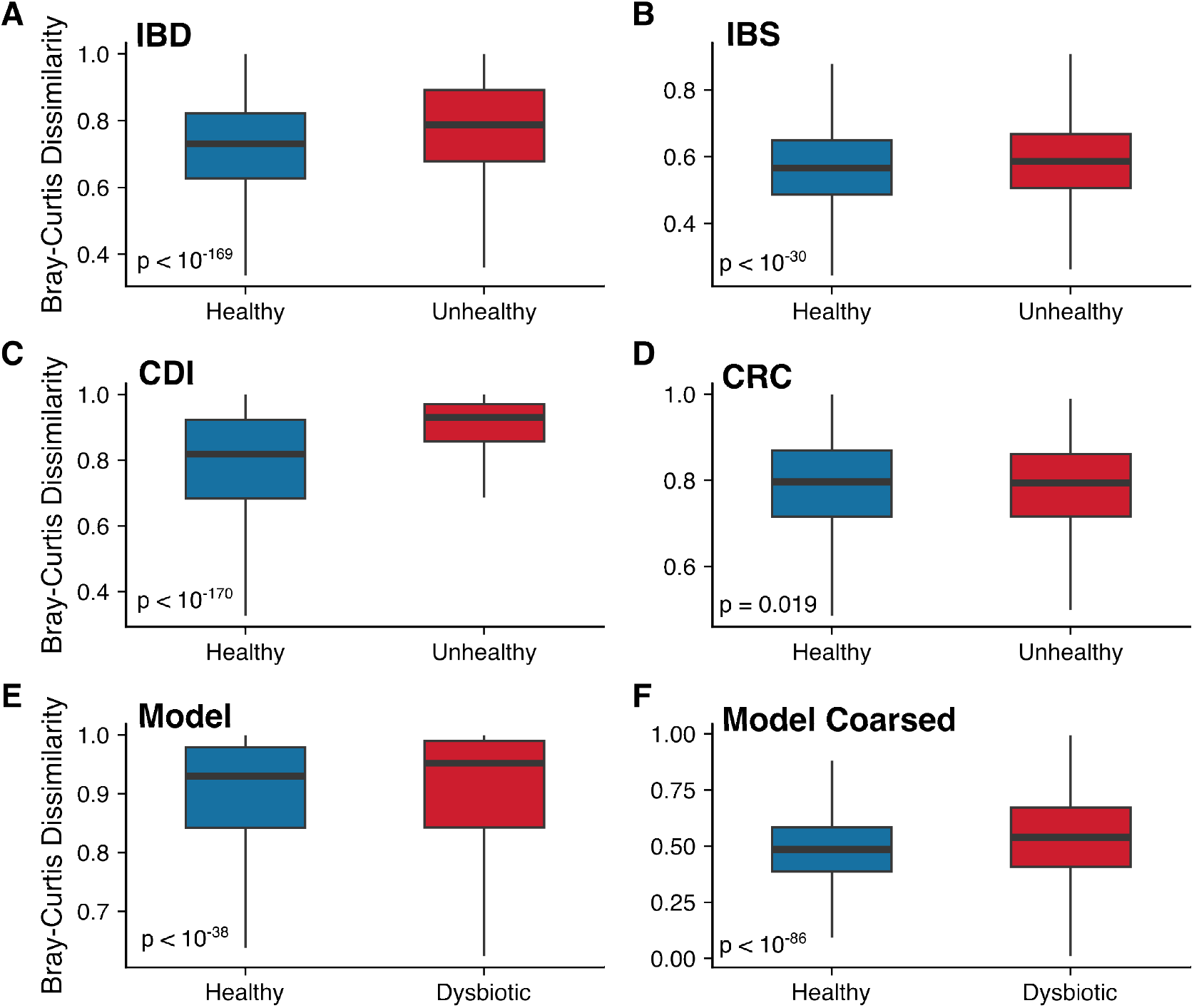
The Anna Karenina Principle holds across different diseases, but not all, and in the model. Beta diversity calculated with the Bray-Curtis dissimilarity of healthy and diseased states for (**A**) IBD, (**B**) IBS, (**C**) CDI, (**D**) CRC, as well as for healthy and collapsed states in the model (**E**) and in the model with coarse-grained pathways (**F**) (see Methods (*86*)). Except for CRC, all diseases and the model follow the Anna Karenina Principle, which means that healthy communities are more similar to each other than dysbiotic ones. The presence of exceptions and the small magnitude of the increases in dissimilarity do not allow for strong conclusions from this observation (see (*109*)).

**Figure S7:**
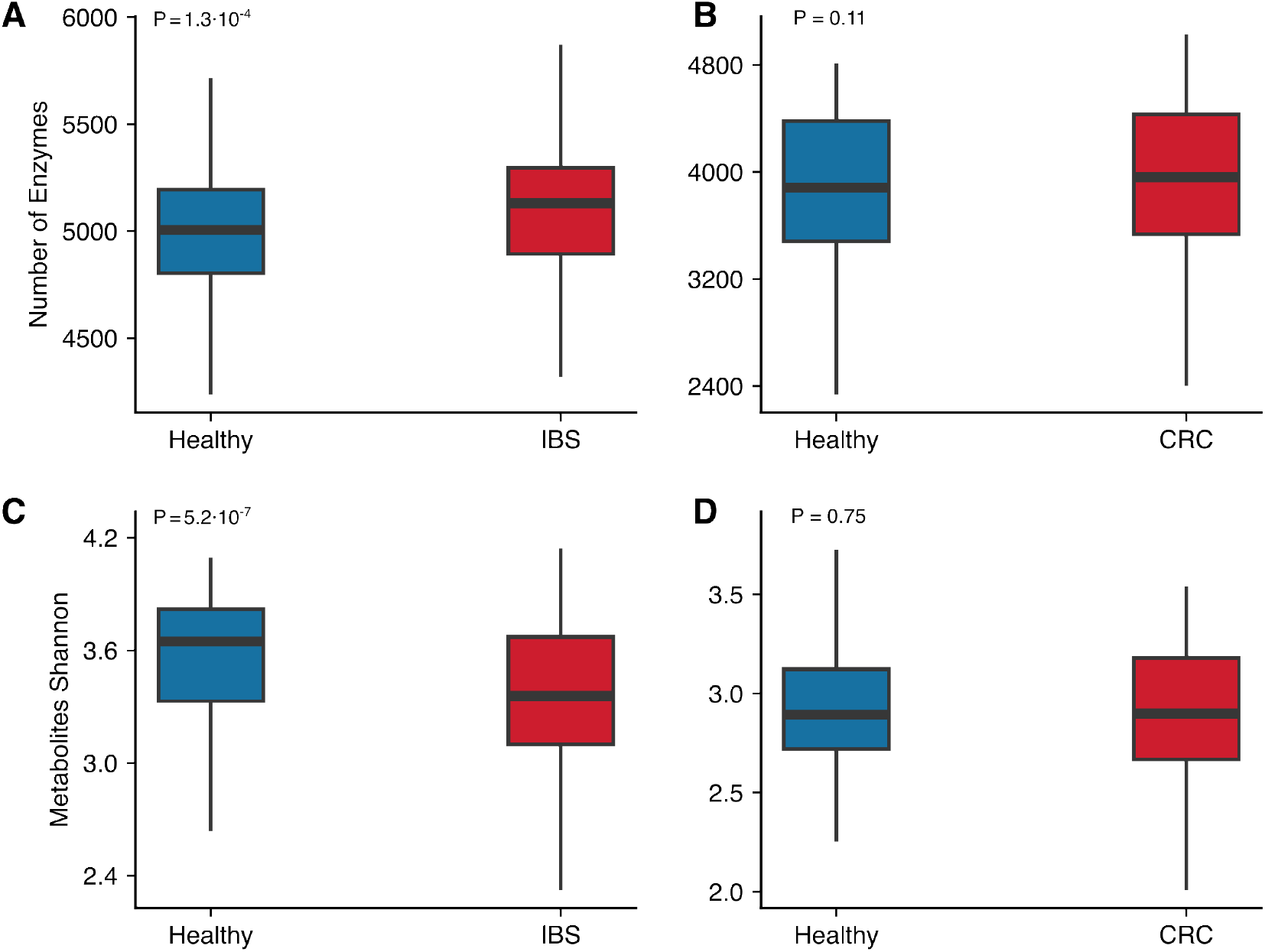
Analyzed biomarkers are not consistently reduced across diseases. (**A-B**) Number of enzymes in Healthy vs. IBS (**A**) and Healthy vs. CRC (**B**). (**C-D**) Shannon index of secreted metabolites in Healthy vs. IBS (**C**) and Healthy vs. CRC (**D**). The inconsistencies observed across diseases indicate that these biomarkers alone are not reliable indicators of dysbiosis.

**Figure S8:**
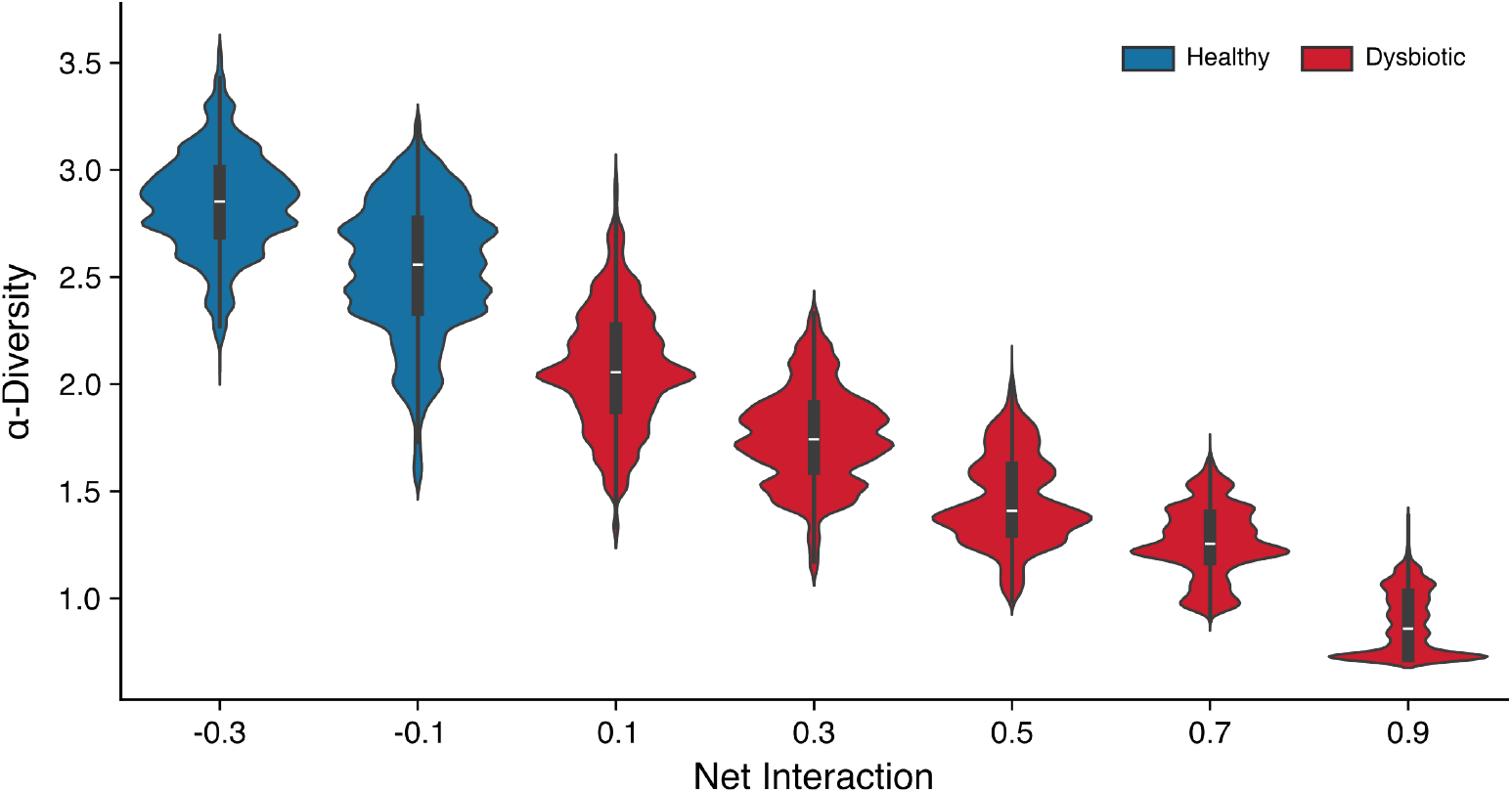
Diversity exhibits high variability across net interaction values. The mean and median of *α*-diversity for a given value of the net interaction shows a negative correlation between the Shannon index and *ρ*. Diversity is, however, widely distributed, which suggests that diversity alone is not a reliable indicator of the underlying interaction structure. For example, for the data in the figure, a value of *α*-diversity of 1.5 could be observed for any network with net interaction in [−0.1, 0.7]. All together supports the claim that a switch to dysbiotic (i.e. to a cooperative network) typically but not always correlates with a decrease in diversity.

**Figure S9:**
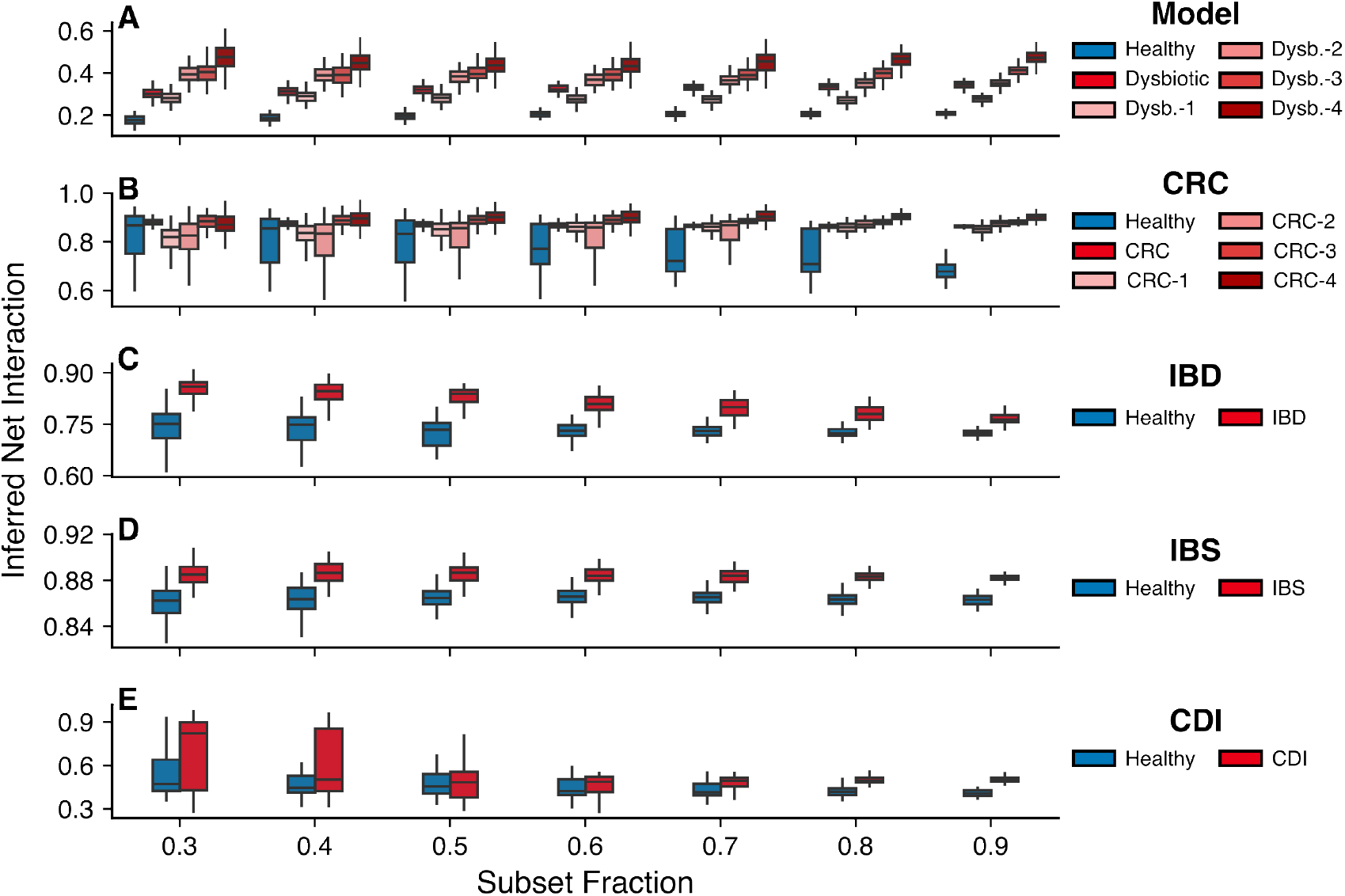
The inferred net interaction is robust under bootstrapping. Inferred net interaction calculated for different subsampling fractions of the original dataset for the model (**A**), CRC (**B**), IBD (**C**), IBS (**D**), and CDI (**E**). The inferred net interaction reliably captures the qualitative trends of the true underlying net interaction, *ρ*, in the model across all subsampled datasets. Healthy (healthy) states consistently show lower inferred net interaction compared to diseased (dysbiotic) states across all datasets. Additionally, for CRC, the inferred net interaction correlates with disease progression, with more advanced disease stages displaying systematically higher inferred net interaction across all subsampled sets. All results are highly statistically significant (500 bootstraping-generated samples)

**Figure S10:**
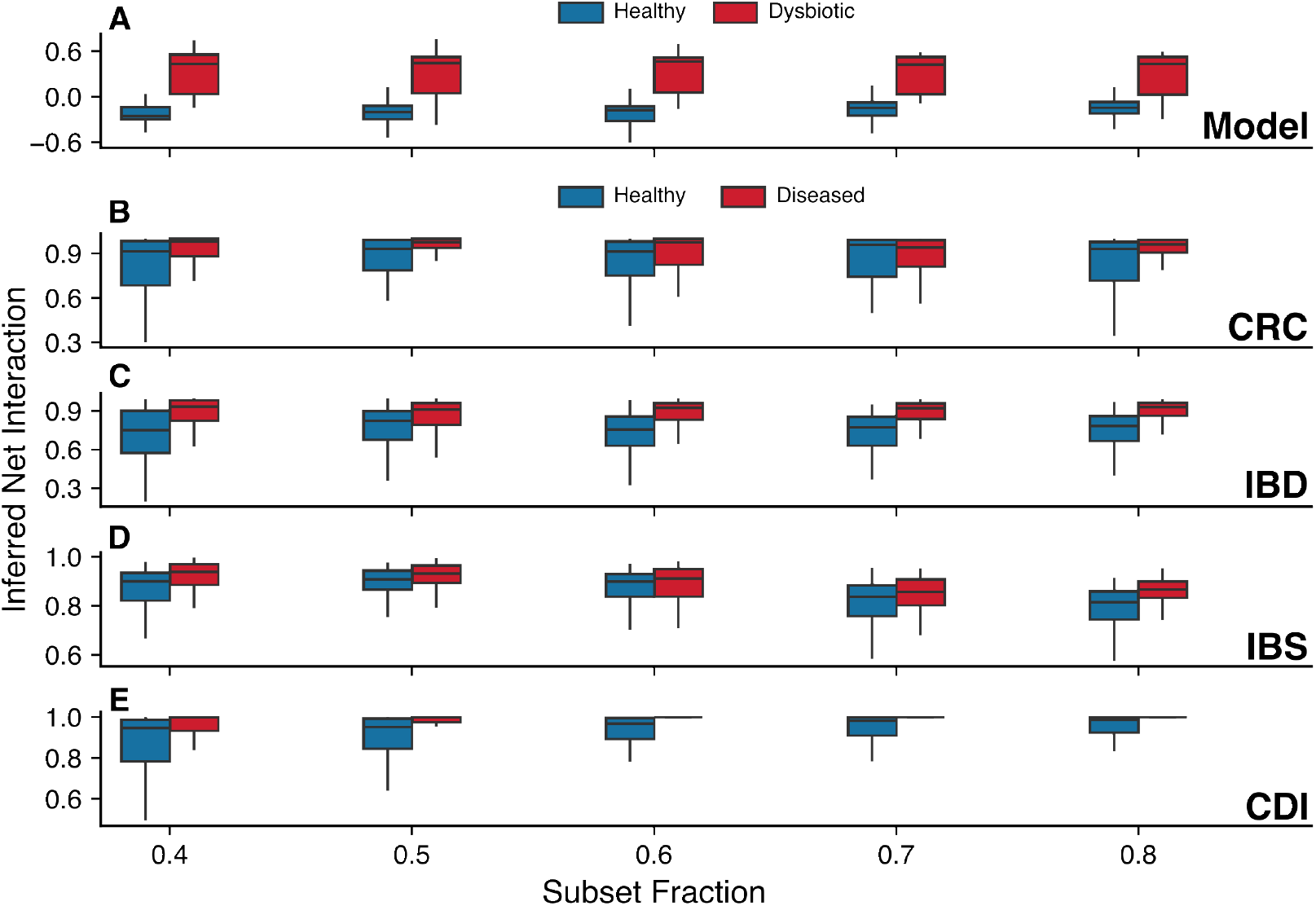
Inferred net interaction using *BEEM-Static*. Inferred net interaction estimated with the *BEEM-Static* algorithm (*105*) (see (*86*)) for different subsampling fractions of the original dataset for the model (**A**), CRC (**B**), IBD (**C**), IBS (**D**), and CDI (**E**). Across all cases, dysbiotic states in the model and disease states in real datasets show a consistently higher inferred net interaction, suggesting a shift toward more cooperative microbial communities compared to healthy states and healthy controls, respectively. All results are highly statistically significant (500 bootstrapinggenerated samples).

**Figure S11:**
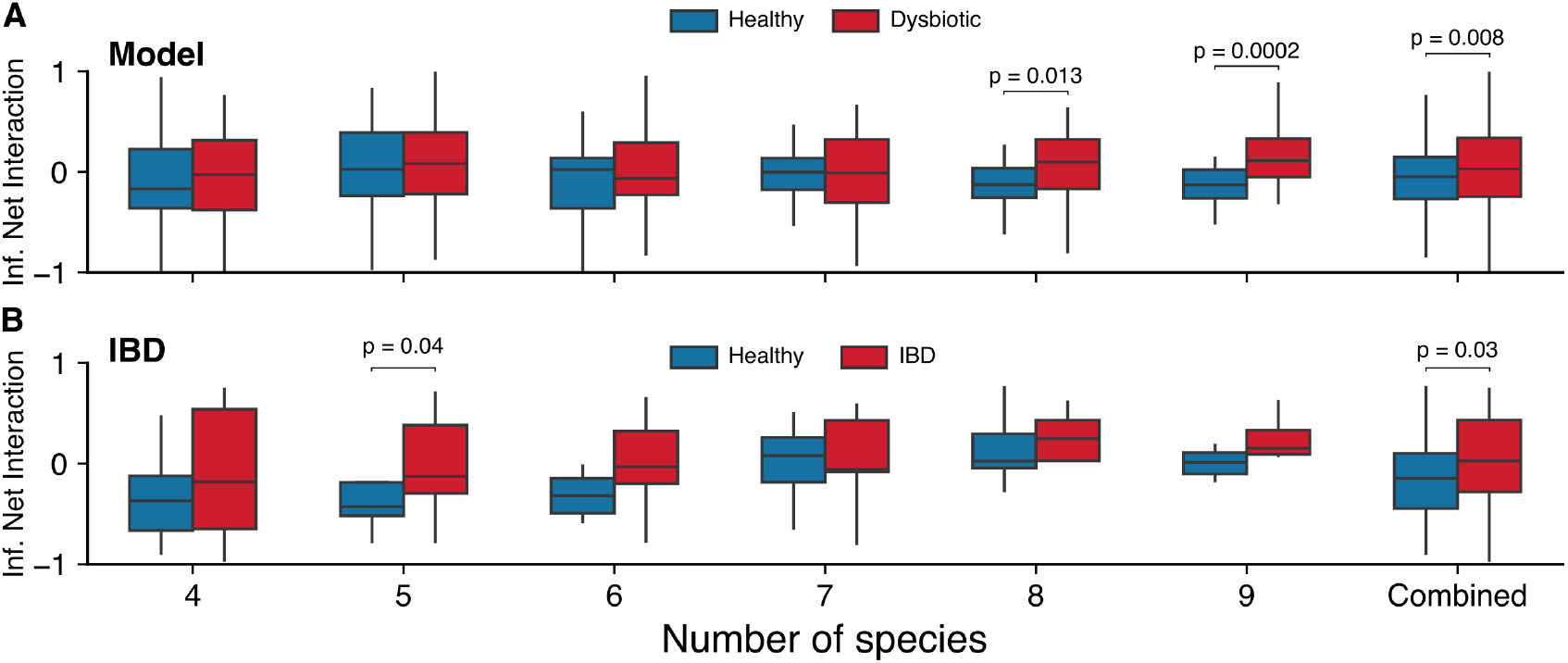
Inferred net interaction using *LIMITS*. Inferred net interaction computed with the *LIMITS* algorithm (*106*) (see (*86*)) for different numbers of species in the model (**A**) and for the IBD dataset (**B**). In both cases, dysbiotic states in the model and diseased states in IBD patients show a higher inferred net interaction compared to their respective healthy counterparts, indicating a shift towards a more cooperative microbial network.

**Figure S12:**
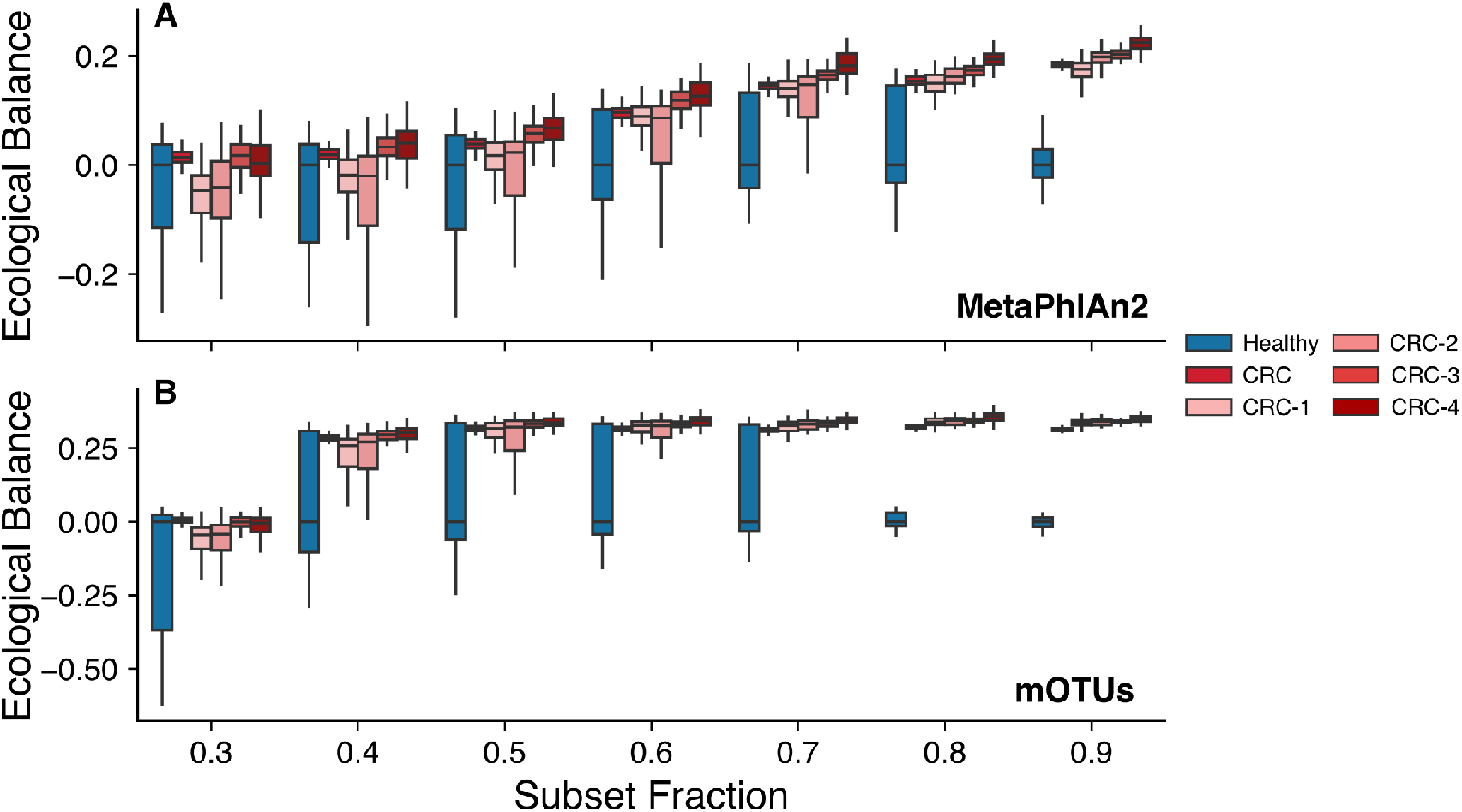
The ecological network balance index (ENBI) is robust across different taxonomic profiling methodologies. ENBI calculated for different subsampling fractions of the original CRC datasets using two distinct taxonomic profiling methodologies: *MetaPhlAn2* (*122*) (**A**) and *mOTUs* (*123*) (**B**). In both cases, disease states consistently show a positive ENBI value, indicating a shift toward more cooperative microbial communities compared to controls. Furthermore, ENBI correlates with disease progression across all subsampled datasets, reinforcing its robustness as a biomarker irrespective of the taxonomic profiling method used. All results are highly statistically significant (500 bootstraping-generated samples).

## Notes

### Competing Interest Statement

The authors have declared no competing interest.

### Summary of Updates

The only changes made have been to change the authors names. - Juan Bonachela to Juan A Bonachela, - SImon Levin to Simon A Levin - Maria Gloria Dominguez Bello to Maria Gloria Dominguez-Bello - Martin Blaser to Martin J. Blaser

